# Mapping the coupling between tract reachability and cortical geometry of the human brain

**DOI:** 10.1101/2025.03.31.646498

**Authors:** Deying Li, Andrew Zalesky, Yufan Wang, Haiyan Wang, Liang Ma, Luqi Cheng, Tobias Banaschewski, Gareth J. Barker, Arun L.W. Bokde, Rüdiger Brühl, Sylvane Desrivières, Herta Flor, Hugh Garavan, Penny Gowland, Antoine Grigis, Andreas Heinz, Herve Lemaitre, Jean-Luc Martinot, Marie-Laure Paillère Martinot, Eric Artiges, Frauke Nees, Dimitri Papadopoulos Orfanos, Luise Poustka, Michael N. Smolka, Nilakshi Vaidya, Henrik Walter, Robert Whelan, Gunter Schumann, Tianye Jia, Congying Chu, Lingzhong Fan, IMAGEN Consortium

## Abstract

The study of cortical geometry and connectivity is prevalent in research on the human brain. However, these two aspects of brain structure are usually examined separately, leaving the essential connections between the brain’s folding patterns and white matter connectivity unexplored. In this study, we aimed to elucidate fundamental links between cortical geometry and white matter tract connectivity. We developed the concept of tract-geometry coupling (TGC) by optimizing the alignment between tract connectivity to the cortex and multiscale cortical geometry. Specifically, spectral analyses of the cortical surface yielded a set of geometrical eigenmodes, which were then used to explain the locations on the cortical surface reached by specific white matter tracts, referred to as tract reachability. In two independent datasets, we confirmed that tract reachability was well characterized by cortical geometry. We further observed that TGC had high test-retest ability and was specific to each individual. Interestingly, low-frequency TGC was found to be heritable and more informative than the high-frequency components in behavior prediction. Finally, we found that TGC could reproduce task-evoked cortical activation patterns. Collectively, our study provides a new approach to mapping coupling between cortical geometry and connectivity, highlighting how these two aspects jointly shape the connected brain.

## Introduction

White matter tracts in the human brain form anatomical connections between different regions, collectively constituting the human connectome ^1,2^. These tracts contribute to information integration and activity coordination across the cerebral cortex, facilitating complex cognitive functions ^3,4^. However, few studies have sought to elucidate the patterns by which white matter tracts connect different parts of the cerebral cortex, i.e., the tract reachability ^2,5,6^, represented by the probability of a tract terminating on the cortical surface. While the macroscale human connectome provides a detailed map of region-to-region connectivity, it does not fully account for the spatial and morphological characteristics of the regions innervated by each tract ^7,8^. Furthermore, as posited by tension-based theory, mechanical tension generated by axons interacts with the gyri-sulci formation during morphogenesis ^9,10^, implying that white matter tracts might interact with cortical geometry and its development. Thus, establishing a link between these two aspects is critical to understanding brain maturation and aging across the lifespan.

Cortical gyrification begins during embryonic development and yields complex cortical geometry to maintain a large surface area relative to a limited cranial volume ^11,12^, accompanied by significant white matter changes ^13^. Evidence has shown that developmental increases in cortical gyrification are associated with disproportionate cortical expansion relative to subcortical structures^14^. White matter undergoes significant concurrent changes, including subplate thickening ^15^, increasing complexity of cortico-cortical fibers, and the emergence of short association fibers around the gyri ^13^. Several recent studies have demonstrated that cortical geometry is linked to brain functions and behaviors ^16,17^, and its abnormal changes could be partly explained by axonal fibers ^18,19^. These findings underscore the interactive coupling between cortical geometry and white matter tracts, implying a dynamic interplay that is crucial for normal brain development and function. However, the precise nature by which cortical geometry and connectivity are coupled remains unknown, and measures are needed to quantify the extent of this coupling.

In this study, we aimed to delineate the coupling between white matter tracts and cortical geometry in the human brain using high-quality multimodal MRI data. We extracted the spatial extent of white matter tracts utilizing diffusion-weighted MRI and characterized the spatial expanse of cortical surface locations intersected by each tract through a probabilistic distribution termed tract reachability. Building on recent work by Pang and colleagues, who employed a multiscale description of cortical geometry to elucidate patterns of cortical connectivity and activation during both task-evoked and resting-state conditions ^20–24^, we integrated cortical geometry into our analysis of complex cortical characteristics. Specifically, we decomposed the cortical surface into geometric eigenmodes, which served as a basis for interpreting tract reachability. This novel approach of correlating tract reachability with cortical eigenmodes is termed tract-geometry coupling (TGC), offering a fresh perspective on the interplay between white matter tracts and cortical structure. An overview of analysis is shown in Figure 1.

**Figure 1.**
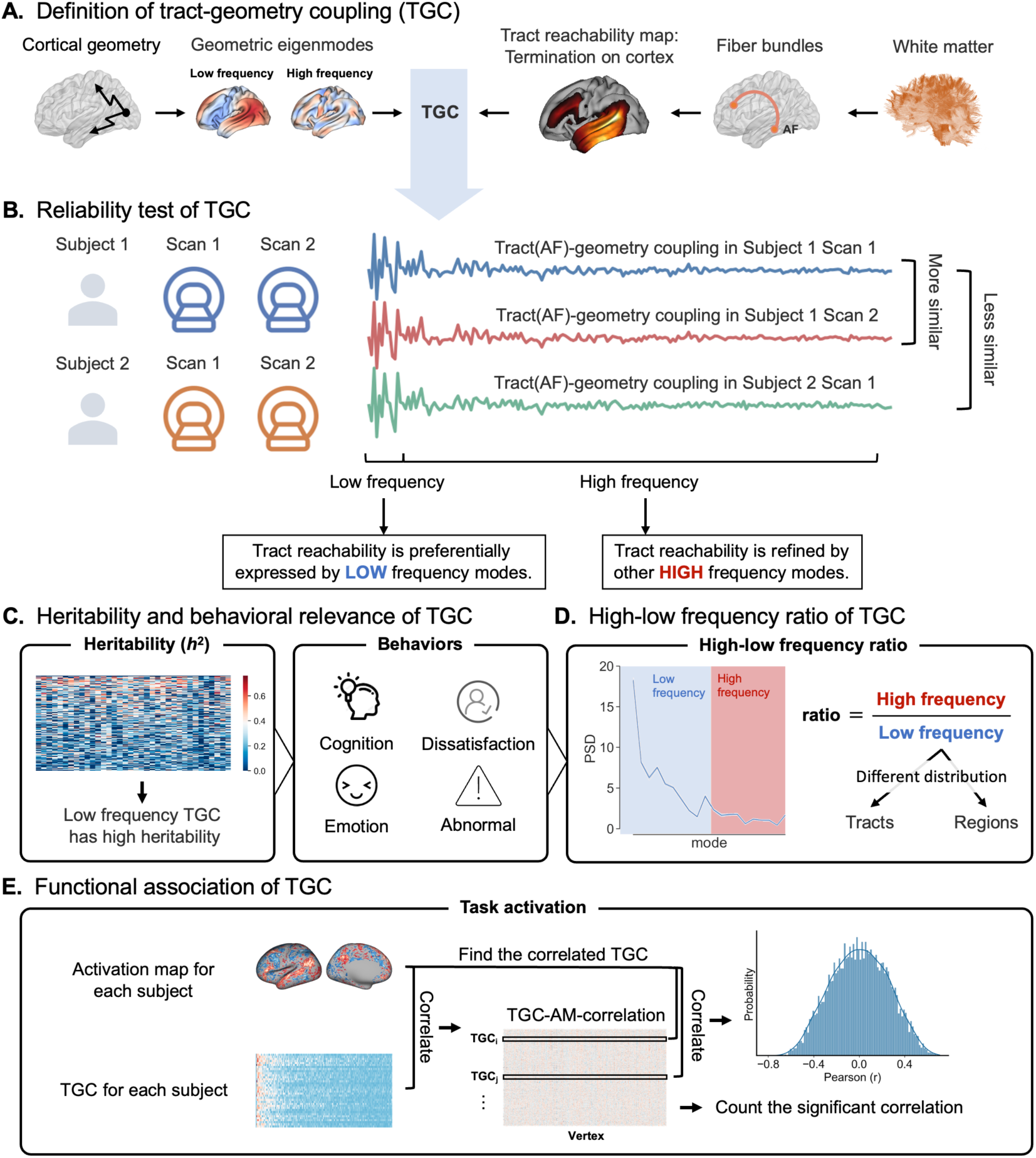
Overview of the study. **(A)** Definition of tract-geometry coupling (TGC). We built the TGC to measure the potential relationship between cortical geometry and tract reachability. **(B)** Reliability test of TGC. We showed that TGC was stable across subjects and captured individual-specific traits. **(C)** Heritability and behavioral relevance of TGC. We assessed the heritability of the TGC and its behavioral predictability. **(D)** High-low frequency ratio of TGC and its distribution across regions and tracts. **(E)** Functional association of TGC. We used task activation maps to explore the functional relationship between TGC scores.

Our findings reveal that TGC is a robust measure, exhibiting heritability and predictive power over individual behavioral variation, with low-frequency eigenmodes offering enhanced explanatory potential. The quantitative assessment of the difference between high- and low-frequency TGC, i.e., high-low frequency ratio, highlighted a pronounced coupling between association tracts and high-frequency eigenmodes, with distinct patterns observed across various functional networks. Moreover, a significant correlation was identified between individual TGC profiles and brain activation maps. In conclusion, our TGC measure not only delineates the intricate relationship between cortical geometry and connectivity but also signifies a step forward in enhancing the accuracy of brain mapping techniques.

## Results

### Linking cortical geometry and tract reachability

We established a new measure to index the relationships between cortical geometry and the reachability of white matter tracts. A population-averaged template of the cortical surface was used to map cortical eigenmodes ^24^ (Figure S1), which were used to reconstruct cortical projection patterns of fiber bundles. The fiber bundles were delineated with a pre-trained deep-learning model, TractSeg ^25^, and their reachability to the cortex was quantified using the connectivity blueprint approach, in which the tract volume was multiplied with a gray matter surface-to-whole-brain tractogram ^26^. The columns of the matrix represented the connectivity distribution pattern of cortical vertices, while the rows provided the cortical projection patterns of the white matter tracts, which is explicitly referred to as the “tract reachability map” in the following text.

We evaluated the accuracy of the geometric eigenmodes in capturing the tract reachability map for the Human Connectome Project (HCP) dataset (*n* = 968) ^27^ and IMAGEN dataset (*n* = 661) ^28^. The tract reachability map of each tract was reconstructed using 200 eigenmodes for each subject using general linear models (GLM) (Figure S2), and reconstruction accuracy was quantified by calculating the Pearson correlation between the empirical and reconstructed tract reachability maps.

Both datasets exhibited high reconstruction accuracy, with all correlation coefficients greater than 0.8, although the results from IMAGEN were slightly lower than those from the HCP dataset (Figure 2A). The results were verified by rotating eigenmaps with BrainSMASH ^29^ and eigenstrapping ^30^ (Figure S3). Empirical and reconstructed tract reachability maps of several example tracts are shown in Figure 2B. The link between tract reachability and cortical geometry elucidated through cortical eigenmode is referred to as the tract-geometry coupling (TGC).

**Figure 2.**
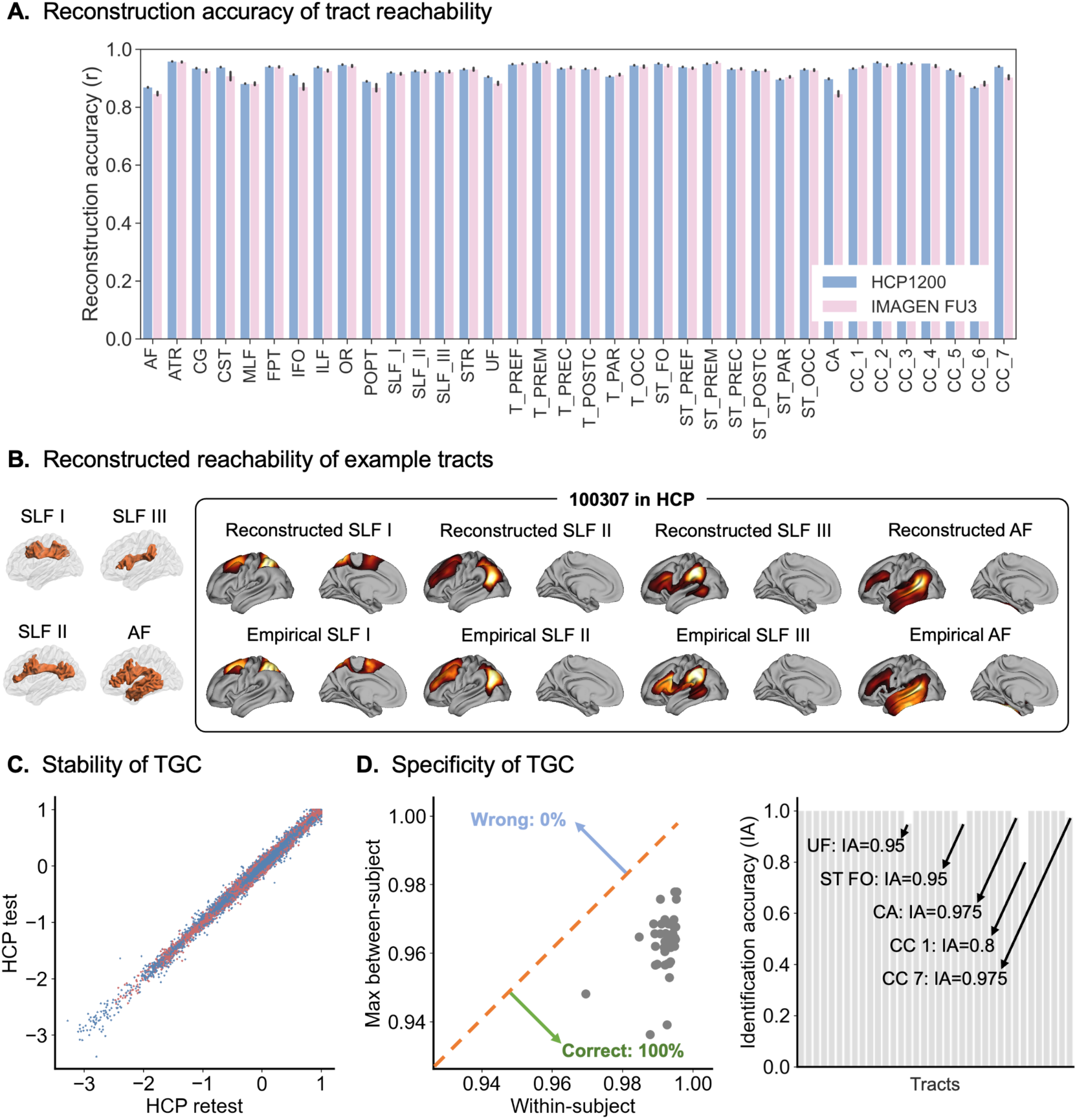
Characterization of TGC. **(A)** Reconstruction accuracy of the tract reachability maps for the HCP and IMAGEN datasets. High accuracy (all correlation coefficients > 0.8) was found for both datasets. **(B)** Empirical and reconstructed reachability maps of several example tracts. **(C)** The TGC is generally stable between the subjects. The similarity of TGC between two scans from the same subject was very high (*r* = 0.99). **(D)** The TGC was able to capture the individual differences across the subjects. The TGC successfully predicted the identities of all subjects (identification accuracy = 100%) in the HCP dataset. SLF I, superior longitudinal fasciculus I; SLF II, superior longitudinal fasciculus II; SLF III, superior longitudinal fasciculus III; AF, arcuate fasciculus; UF, uncinate fascicle; ST_FO, striato-fronto-orbital tract; CA, commissure anterior; CC_1, corpus callosum rostrum part; CC_7, corpus callosum splenium part.

We considered the impact on reconstruction accuracy of including fewer or more eigenmodes. We found that reconstruction accuracy increased with the number of eigenmodes across all tract reachability maps, with *r* = 0.5 achieved using just *N* = 10 modes and *r* > 0.7 using *N* = 50 (Figure S4), and remained high when using individual geometric eigenmodes (Figure S5A). Meanwhile, TGC calculated by individual geometric eigenmodes showed high similarity to TGC calculated by group geometric eigenmodes (Figure S5B), so we used the latter in the further analysis. The reproducible results of the right hemisphere demonstrated the same level of high accuracy (Figure S6). Given that the geometric eigenmodes of each hemisphere were decomposed separately, we did not combine the TGC from the two hemispheres. Therefore, we primarily used the TGC from the left hemisphere as the main results.

### TGC is stable and specific across individuals

To explore the stability of the TGC, we measured the similarity of the TGC between two scans of the same subject using a cohort from the HCP who underwent test and retest scanning over several days (HCP test-retest, *n* = 44). We found that the correlation between the two scans was very high (*r* = 0.99; Figure 2C).

We also investigated whether the TGC could capture individual differences between subjects. We performed the analysis by measuring the similarity between each pair of subjects from two scans in the HCP test-retest dataset ^31^. For example, for an individual from the first scan, the Pearson correlation coefficient was calculated between this individual and all subjects from the second scan. If the coefficient for a given subject was greater than the maximum coefficient of all the other subjects, the identity of this subject was predicted. The results showed that the TGC could predict the identities of all the subjects (identification accuracy [IA] = 100%), indicating its ability to reflect individual differences (Figure 2D, left). We repeated the same procedure using the TGC of each tract separately and found similar results, except for five white matter tracts (UF: IA = 95%, ST_FO: IA = 95%, CA: IA = 97.5%, CC_1: IA = 80%, CC_7: IA = 97.5%; Figure 2D, right).

### Low-frequency TGC is heritable and associated with behavior

We found a stronger coupling relationship in the low-frequency eigenmodes, while a weaker relationship was observed in the high-frequency eigenmodes (Figure 3A, 3B). This suggests that high-frequency eigenmodes may capture individual differences in the coupling between cortical geometry and connectivity. This motivated us to evaluate TGC heritability using twins from the HCP dataset (*n* = 194, 102 monozygotic and 92 dizygotic pairs). We found that low-frequency eigenmodes showed greater TGC heritability than high-frequency eigenmodes (*r* = −0.74, *p* = 2e-35; Figure 3C, left). This difference was also observed in the association, commissure, and projection tracts (Figure 3C, right).

**Figure 3.**
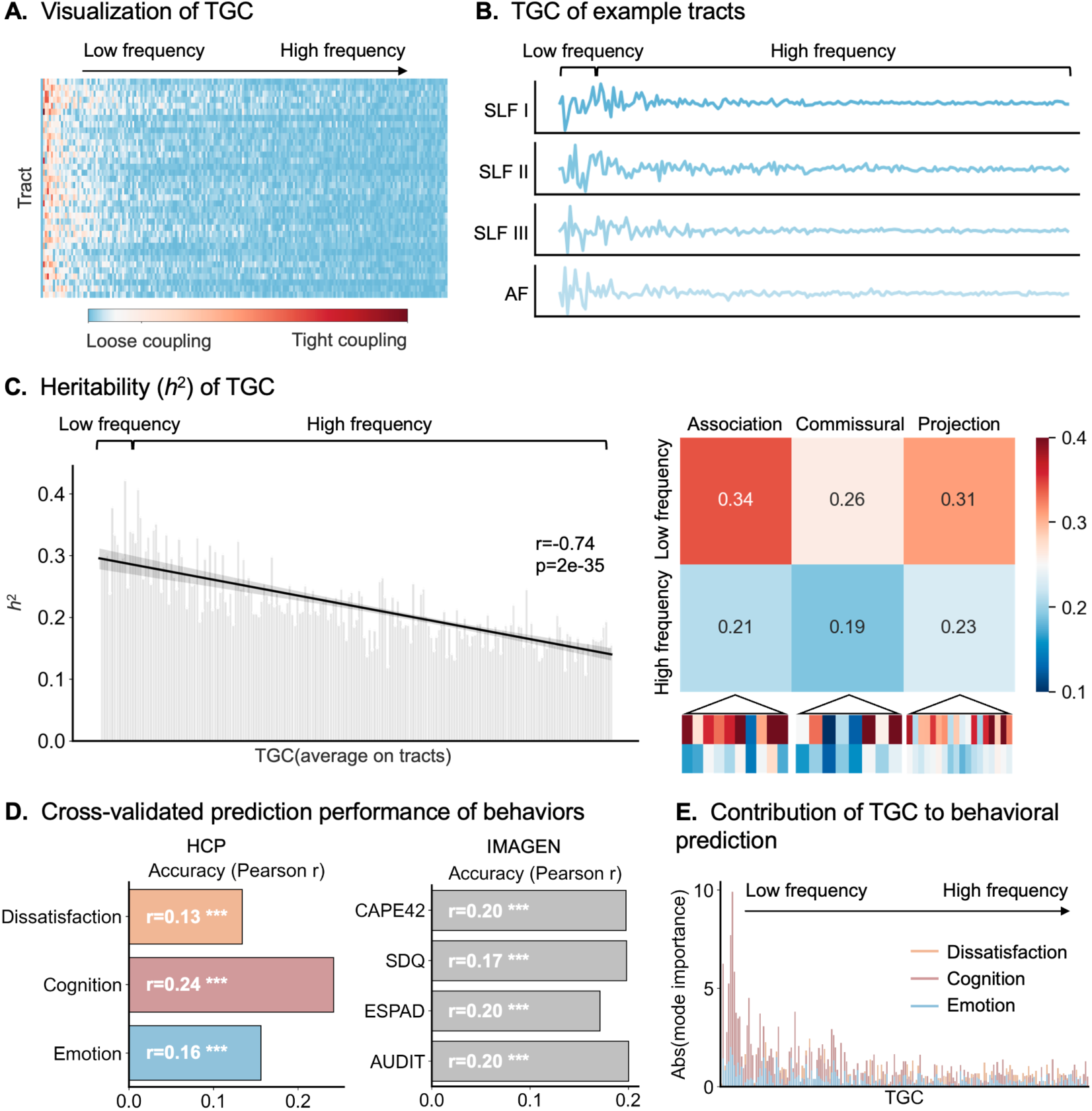
Heritability and behavioral relevance of TGC. **(A)** Visualization of the TGC. A stronger coupling relationship was observed in the low-frequency eigenmodes, while a weaker relationship was found in the high-frequency eigenmodes. **(B)** The TGC for several example tracts. **(C)** Left: Heritability of the TGC. The coupling between tracts and low-frequency eigenmodes showed greater heritability than that with high-frequency eigenmodes. Right: Differences in heritability of the coupling with low-frequency and that with high-frequency eigenmodes were also observed across different tract types. **(D)** The TGC could predict the three components derived from behaviors in the HCP dataset (dissatisfaction: *r* = 0.13, *p*_permutation_ = 0; cognition: *r* = 0.24, *p*_permutation_ = 0; emotion: *r* = 0.16, *p*_permutation_ = 0) and four measures of dysfunctional behaviors in IMAGEN dataset (CAPE42: *r* = 0.20, *p*_permutation_ = 0; SDQ: *r* = 0.17, *p*_permutation_ = 0; FTND: *r* = 0.20, *p*_permutation_ = 0.001; AUDIT: *r* = 0.20, *p*_permutation_ = 0). **(E)** Contribution of TGC to behavioral predictions. The low-frequency TGC showed greater contributions to all three components.

We also found that TGC predicted individual variation in several behaviors. Kernel ridge regression (KRR) was used to predict the three components derived from 58 behavioral scores in the HCP dataset ^32^. We found that the behavior could be well predicted (dissatisfaction: *r* = 0.13, *p*_permutation_ = 0; cognition: *r* = 0.24, *p*_permutation_ = 0; emotion: *r* = 0.16, *p*_permutation_ = 0; Figure 3D, left, Figure S7, Table S1). Feature importance was calculated for each component score using the Haufe transformation ^33^, revealing that low-frequency TGCs were the most informative to behavior prediction (Figure 3E).

We also showed that TGC could predict high-risk behaviors using KRR on the IMAGEN data. The results again showed its ability to reflect functionally abnormal behaviors (CAPE: *r* = 0.20, *p*_permutation_ = 0; SDQ: *r* = 0.20, *p*_permutation_ = 0; FTND: *r* = 0.17, *p*_permutation_ = 0.001; AUDIT: *r* = 0.20, *p*_permutation_ = 0; Figure 3D, right, Figure S7, Table S1) and psychopathology symptoms (depband: *r* = 0.13, *p*_permutation_ = 0.25; spphband: *r* = 0.22, *p*_permutation_ = 0; dcgena: *r* = 0.23, *p*_permutation_ = 0.002; ocdband: *r* = 0.19, *p*_permutation_ = 0.006; eatband: *r* = 0.19, *p*_permutation_ = 0; ptsdband: *r* = 0.15, *p*_permutation_ = 0.014; Figure S8).

Tract reachability was also used to predict behavior and its performance was compared to those using TGC. Both TGC and tract reachability could predict cognitive behaviors in the IMAGEN dataset and TGC has a better performance (Figure S9). Behaviors in the HCP and IMAGEN datasets can also be significantly predicted using TGC in the right hemisphere (Table S1), highlighting the individual variability of TGC in the right hemisphere and its functional associations.

### A high-low frequency ratio shows functional relevance

Given the finding of a stronger coupling relationship between white matter tracts and low-frequency eigenmodes than high-frequency eigenmodes, we sought to provide a quantitative measure for this difference, i.e., the high-low frequency ratio, modified from Preti et al. ^34^. We decomposed the tract reachability maps into two components: one that is closely coupled with geometry, *y^L^*, represented by low-frequency eigenmodes, and another that is less closely coupled, *y^H^*, represented by high-frequency eigenmodes (Figure S10). The ratio between the norm of *y^H^* and *y^L^* across tracts was utilized as the measure of tract-geometry coupling for a specific tract. We calculated the high-low frequency ratio for each tract and assessed the difference across association, commissure, and projection tracts. We found a significant difference between the association and projection tracts (HCP: *t* = 3.50, *p* < 0.005; IMAGEN-FU3: *t* = 2.92, *p* < 0.01; Table S2).

We averaged the high-low frequency ratio map across tracts, resulting in an average map (Figure 4A). We assigned the map to the multimodal parcellation scheme ^35^ and found that the regions showing a high high-low frequency ratio consisted of the dorsolateral prefrontal cortex, inferior parietal lobule, rostroventral insula, and cingulate cortex (Figure 4A), which were located in multiple demand network ^36^ and default mode network ^37^. Similar results were replicated in FU3 cohort from the IMAGEN datasets (*r* = 0.85, *p* = 4e-30; Figure 4B) and using another atlas, the Brainnetome parcellation ^38^ (Figure S11).

**Figure 4.**
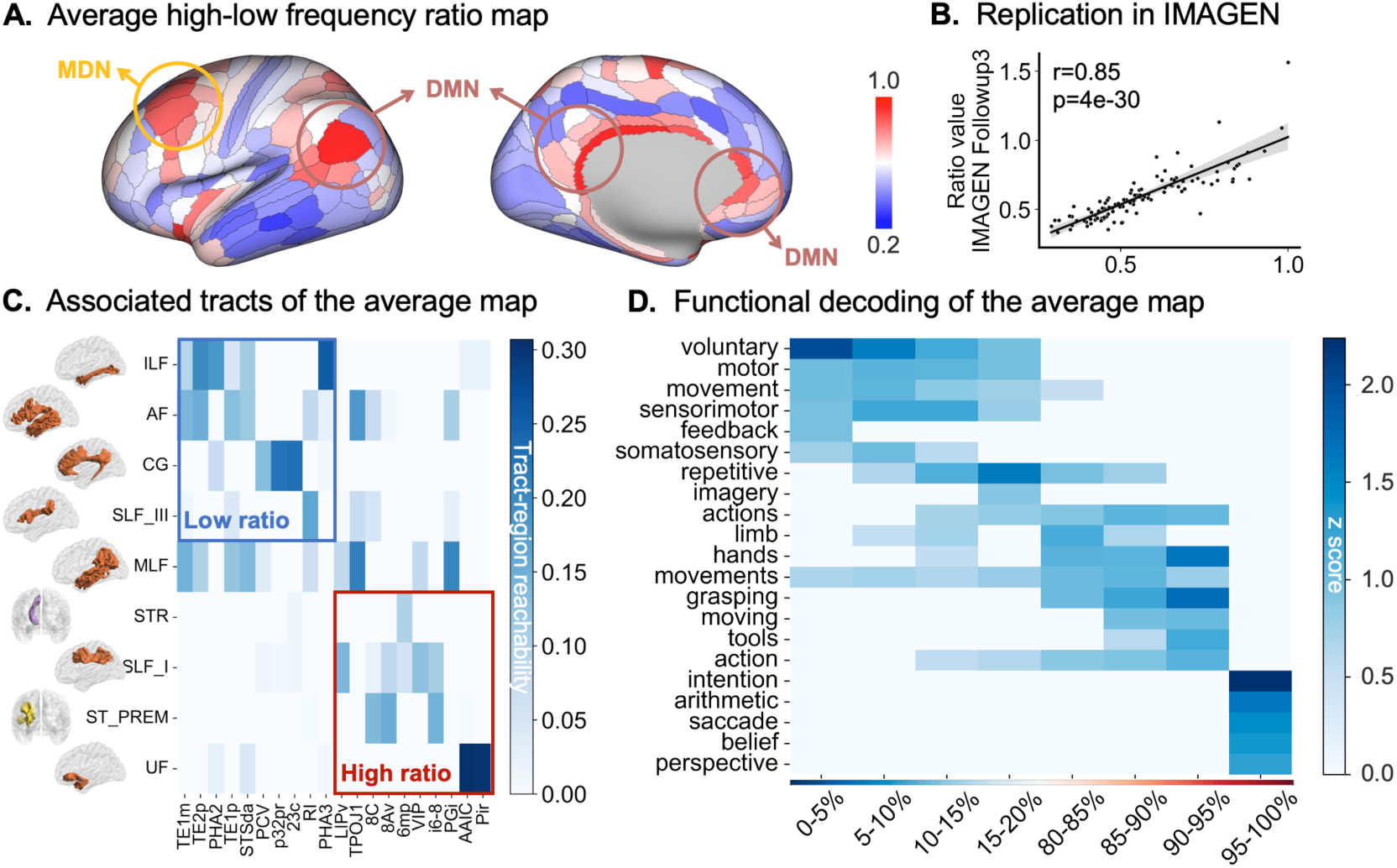
High-low frequency ratio of TGC. **(A)** The average high-low frequency ratio map across tracts. Regions with a higher high-low frequency ratio are located in the dorsolateral prefrontal cortex, inferior parietal lobule, rostroventral insula, and cingulate cortex, which are part of the multiple demand network and default mode network. **(B)** Replication using the FU3 cohort from the IMAGEN datasets (*r* = 0.85, *p* = 4e-30). **(C)** Associated tracts for regions with high and low ratios. **(D)** Functional decoding of the average high-low frequency ratio map.

When investigating the specific white matter tracts these subregions were connected with, we found that low-ratio regions were associated with the ILF, AF, CG, and SLF_III, while high ratio regions were associated with the STR, SLF_I, ST_PREM, and UF (Figure 4C). As the coupling between tracts and low-frequency eigenmodes was more influenced by genetic factors, we hypothesized that individualized and local information was more prevalent in the coupling with high-frequency eigenmodes. This might be reflected in the functional association of the high-low frequency ratio. Functional decoding using NeuroSynth ^39^ showed that regions with a high ratio were related to “intention,” “arithmetic,” “saccade,” and “belief,” whereas regions with a low ratio were involved in “voluntary,” “motor,” “movement,” and “sensorimotor” (Figure 4D).

### TGC is associated with task-evoked activation

We next investigated whether TGC is associated with task-evoked cortical activation maps derived from fMRI. We used task activation maps from 47 contrasts in the HCP data. For each contrast, we correlated each TGC with each cortical vertex across subjects, resulting in a correlation matrix, which was then correlated with the activation map, yielding a correlation for each activation map. We found that the maximum correlation between the TGC and the activation map (TGC-AM-correlation) closely resembled the activation map itself (Figure 5A, Figure S12). For example, a stronger correlation was found in the posterior part of the inferior parietal lobule, which is also the peak activation location of the contrast map for Language Story-Math (Figure 5A, left), which suggests that the variation trend of TGC across the population corresponds to the trend of activation intensity across the population, especially at significantly activated vertex. This observation was further validated by calculating the relationship between the activation value of an example vertex and the TGC across subjects (Figure 5B). These results indicate that the association between TGC and the activation map is continuously and uniformly distributed across subjects, instead of coincidental, rather than being driven by a few individuals with exceptionally high TGC and activation values.

**Figure 5.**
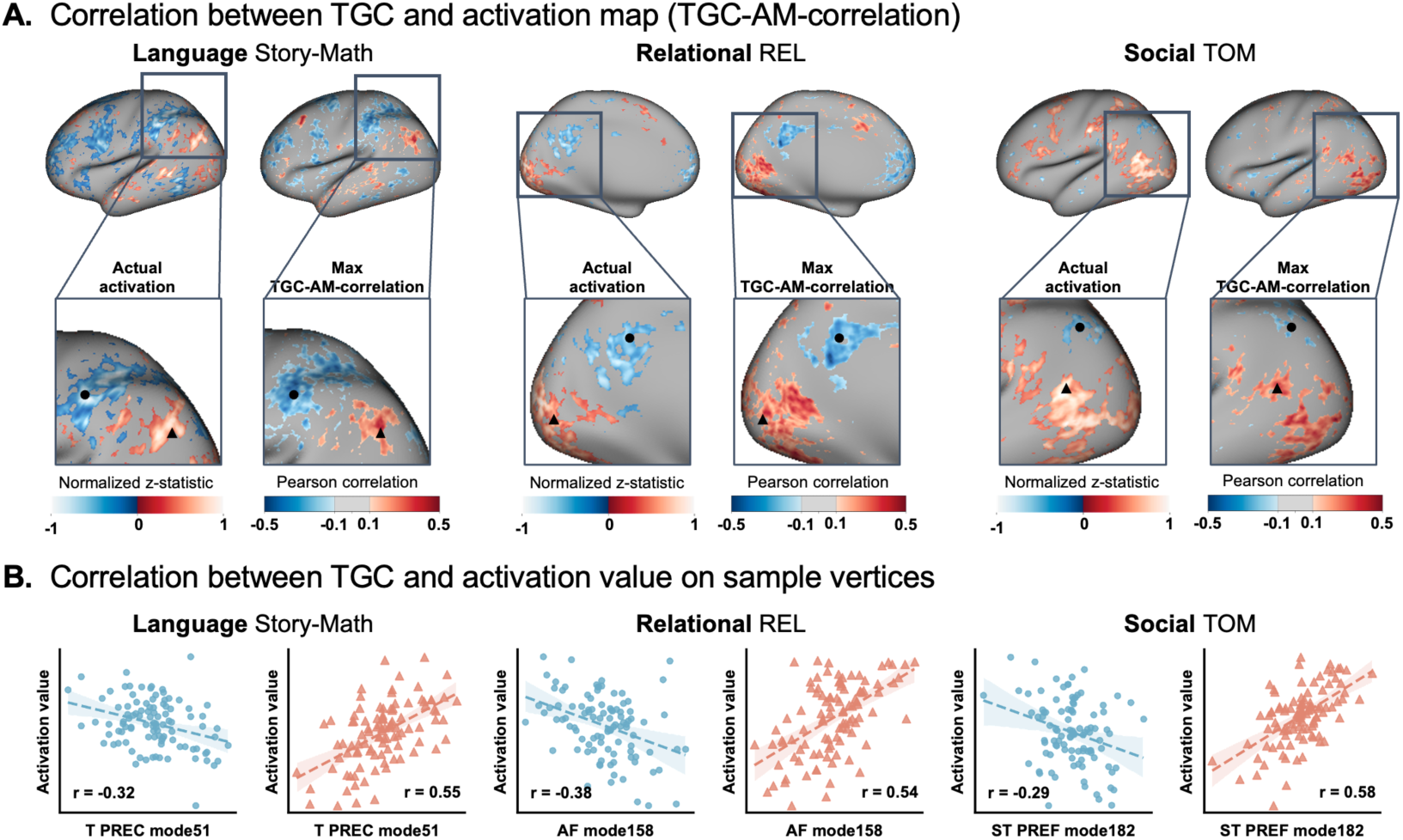
TGC represented functional association. **(A)** The maximum correlation between TGC and the activation map (TGC-AM-correlation) showed a similar pattern to the actual activation map. For example, a stronger correlation was observed in the posterior part of the inferior parietal lobule, which is also the peak activation location of the contrast map for Language Story-Math. **(B)** The activation value at a specific vertex showed a significant correlation with TGC across subjects. The vertex indicated by the black circle in (A) is negatively correlated with TGC, while the vertex indicated by the black triangle in (A) is positively correlated with TGC.

We then quantified the significance of the correlation by first performing multiple comparisons on the correlation matrix described above and counting the number of vertices showing significant correlation values for each TGC. The similarity between activation maps for contrasts within the same behavioral domain was high (Figure S13), indicating that TGC captures functional information relevant to task activation domains.

## Discussion

In this study, we unveiled new insights into the relationship between cortical geometry and connectivity using a measure called tract-geometry coupling (TGC). We found that TGC is a stable measure that is heritable and highly unique to individuals. TGC also explains task-evoked cortical activity and individual variability in behaviors. Based on eigenmodes spanning a spectrum of frequencies, we observed that TGC for high-frequency eigenmodes and association tracts was less heritable than for low-frequency counterparts. This also reflects the greater level of coupling between high-frequency eigenmodes and association tracts. Finally, we showed that the TGC demonstrated a significant correlation with brain activation.

Previous studies have focused on region-to-region connectivity but have overlooked the influence of white matter on these regions, which are physically embedded in three-dimensional space and manifest complex geometric shapes. This gap motivates us to consider the mutual constraints between cortical geometry and white matter pathways, aiding our understanding of how the brain’s architecture gives rise to cognition and behavior ^3,24^. The formation of geometric patterns in gray matter is supported by the underlying white matter skeleton, which also constrains the direction and range of axonal growth. This process begins during the fetal period ^40^ and continues to evolve through childhood and adolescence ^41,42^. This phenomenon could be explained by synaptic construction, increased axonal diameter, and myelination at the microstructural level, as indicated by physical-mechanical models, lesion studies, and molecular experiments ^43,44^. However, studying associations of a single brain region has its limitations. Therefore, we proposed to characterize the coupling relationship between tract connectivity and cortical geometry from a global and macroscale perspective. Tract reachability, characterized using connectivity blueprints that help mitigate bias to some extent when fiber bundle enters the gray matter, provides a comprehensive description of tract-to-cortex connectivity ^26,45^. We provided clear evidence that tract reachability was preferentially expressed using low-frequency, smooth spatial patterns obtained from the harmonic decomposition of the cortical geometry. In addition to the highly repeatable observations of this coupling across different cohorts, we demonstrated both stability and uniqueness in each subject, as well as individual predictability for cognitive and high-risk abnormal behaviors. These findings suggested that tract-geometry coupling is a reliable indicator of the relationship between cortical geometry and white matter connectivity.

From the visualization of various cortical geometric eigenmodes, the low-frequency components showed typical anterior-posterior and dorsal-ventral patterns, with clear separations across different brain lobes. As frequency increased, more refined brain regions or networks, including the insula, default mode network, and motor areas, became increasingly distinct. Additionally, we found greater heritability in the coupling between low-frequency eigenmodes and white matter tracts compared to high-frequency eigenmodes. This could be explained by the morphogens that guide axons along the dorsal-ventral, rostral-caudal, and lateral-medial axes during early development, which aligns with the low-frequency geometric eigenmodes, contributing to the functional arealization of the cerebral cortex in adults ^46^. This process further leads to varying functional significance across different cortical areas. In addition, greater heritability was found in projection tracts, consistent with their conserved roles across subjects during phylogenesis and ontogenesis ^47,48^. In contrast, individual differences in cortical structure and functions were more pronounced in the high-frequency components, which influence the connectional patterns of association tracts ^2,49^.

In a recent study, Pang et al. constructed a more compact and accurate macroscopic expression of brain function based on the geometric basis of the cerebral cortex ^24^ and captured the fundamental anatomical constraints of brain dynamics. White matter networks support the realization of brain functions and promote effective and coordinated information transmission, including both spontaneous and evoked brain functional activities ^3,50,51^ across regions ^52^, while also providing a basis for the functional dynamics of the brain through modern biophysical models and network control theory ^53–56^. In this study, the coupling between cortical geometry and tract reachability across the entire cortex showed greater decoupling in most of the association cortex, particularly in multiple demand network and default mode network which are domain-general in cognitive operations, indicating that high-frequency, short-range and local geometric eigenmodes interacted more strongly with white matter tracts than low-frequency ones. This consistent relationship is also supported by the predictability of TGC for different aspects of brain functional activations and behaviors. That indicates that TGC could serve as a valuable reference for clinical and functional studies, enabling the investigation of how variations in these connections might relate to neurological and psychiatric conditions, thus supporting the development of new diagnostic tools and therapeutic strategies aimed at targeting specific white matter pathways.

Several technical and methodological limitations should be acknowledged. The first and most direct issue is the accuracy of using diffusion MRI to map structural connectivity. Gyral bias caused by the current reconstruction and tractography methods may lead to false positive results, especially when encountering crossing fibers ^57–59^. In addition, dMRI can only evaluate the microstructure of white matter indirectly and is limited in its ability to describe intra-axial characteristics, especially at lower diffusion weights. This makes it difficult to interpret specific indicators related to development as precise microstructural events ^60,61^. In our current work, we quantitatively analyzed the coupling relationship between cortical geometric patterns and white matter tract reachability in two adult brain datasets. However, what is even more interesting is how this coupling relationship is formed and developed during development and evolution. In addition, previous studies have shown that patients with neurodevelopmental disorders exhibit abnormalities in cortical maturation and white matter connectivity ^62,63^. In the future, more attention should be given to studying how this coupling affects these atypical populations.

In summary, we reconstructed white matter tract reachability using geometric eigenmodes and proposed tract-geometry coupling (TGC) to explore the effect of cortical geometric eigenmodes on white matter tract reachability. We demonstrated that TGC is a stable and specific measure and found a distinct heritability between low-frequency and high-frequency eigenmodes and between association and projection tracts. We further demonstrated that association tracts and high-frequency eigenmodes have a stronger coupling relationship and is related to fiber length profiles. Finally, we showed that TGC could predict brain activation patterns. In short, we proposed a new perspective for comprehending the relationship between cortical geometry and white matter tracts, providing insights into the functioning of the human brain.

## Materials and Methods

### Data acquisition

#### HCP dataset

We used a publicly available dataset containing 968 subjects (459 males; mean age, 28.70±3.71; age range, 22-35) provided by the Human Connectome Project (HCP) database ^27^ (http://www.humanconnectome.org/). All the scans and data from the individuals included in the study had passed the HCP quality control and assurance standards.

The scanning procedures and acquisition parameters were detailed in previous publications ^64^. In brief, T1w images were acquired with a 3D MPRAGE sequence on a Siemens 3T Skyra scanner equipped with a 32-channel head coil with the following parameters: TR = 2400 ms, TE = 2.14 ms, flip angle = 8°, FOV = 224×320 mm^2^, voxel size = 0.7 mm isotropic. Diffusion data were acquired using single-shot 2D spin-echo multiband echo planar imaging on a Siemens 3 Tesla Skyra system (TR = 5520 ms, TE = 89.5 ms, flip angle = 78°, FOV = 210×180 mm). These consisted of three shells (b-values = 1000, 2000, and 3000 s/mm^2^), with 90 diffusion directions isotropically distributed among each shell and six b = 0 acquisitions within each shell, with a spatial resolution of 1.25 mm isotropic voxels.

#### IMAGEN dataset

We also included the IMAGEN dataset in our analysis ^28^. IMAGEN is a large-scale longitudinal neuroimaging-genetics cohort study conducted to understand the biological basis of individual variability in psychological and behavioral traits and their relationship to common psychiatric disorders. The participants were recruited from schools in France, the UK, Ireland, and Germany. MRI data were acquired at eight IMAGEN assessment sites with 3T MRI scanners from different manufacturers (Siemens, Philips, GE Healthcare, Bruker). The study involved a thorough neuropsychological, behavioral, clinical, and environmental assessment of each participant. The participants also underwent a biological characterization that included the collection of T1-weighted structural MRI and diffusion MRI data. A more detailed description can be found in the standard operating procedures for the IMAGEN project (https://imagen-europe.com/resources/standard-operating-procedures/). In this investigation, we used T1-weighted structural MRI data, diffusion MRI data, and behavior data from ages 14 (baseline), 19 (follow-up2), and 22 (follow-up3). We only included 661 participants (304 males; mean age, 14.43±0.39 in the baseline, 18.97±0.68 in follow-up2, 22.46±0.62 in follow-up3) who had complete neuroimage data and demographic information, including age, gender, handedness, and acquisition site. The scanning variables were specifically chosen to be compatible with all the scanners. Diffusion imaging protocols were specifically harmonized across sites and scanners for the IMAGEN study, while structural scans were based on the protocols developed by the Alzheimer’s Disease Neuroimaging Initiative (ADNI) (https://adni.loni.usc.edu/data-samples/adni-data/neuroimaging/mri/mri-scanner-protocols/), which are optimized to provide very similar results despite differing scanner capabilities and thus differing acquisition parameters.

### Image preprocessing

#### HCP dataset

The human T1w structural data had been preprocessed following the HCP’s minimal preprocessing pipeline ^64^. In brief, the processing pipeline included imaging alignment to standard volume space using FSL ^65^, automatic anatomical surface reconstruction using FreeSurfer ^66^, and registration to a group average surface template space using the multimodal surface matching (MSM) algorithm ^67^. Human volume data were registered to the Montreal Neurological Institute (MNI) standard space, and surface data were transformed into surface template space (fs_LR).

The diffusion images were processed using FDT (FMRIB’s Diffusion Toolbox) of FSL ^65^. The main steps are normalization of the b0 image intensity across runs and correction for echo-planar imaging (EPI) susceptibility, eddy-current-induced distortions, gradient-nonlinearities, and subject motion. DTIFIT was then used to fit a diffusion tensor model. The probability distributions of the fiber orientation distribution were estimated using Bedpostx ^68,69^.

Next, skull-stripped T1-weighted images for each subject were co-registered to the subject’s b0 images using FSL’s FLIRT algorithm. Then, nonlinear transformations between the T1 image and the MNI structural template were obtained using FSL’s FNIRT. Based on concatenating these, we derived bi-directional transformations between the diffusion and MNI spaces.

#### IMAGEN dataset

For the IMAGEN dataset, processing of the structural T1-weighted images was performed following the HCP’s minimal preprocessing pipeline. The processing pipeline included image alignment to standard volume space using FSL, automatic anatomical surface reconstruction using FreeSurfer, and registration to a group average surface template space. Human volume data were registered to the MNI standard space, and surface data were transformed into surface template space (fs_LR).

The diffusion MRI was also processed using FDT. The main pipeline began with normalizing the b0 intensity, followed by estimating the EPI distortion, eddy current distortions, and subject head motion into a Gaussian process predictor to allow for correction. Then the b0 image was registered to the T1-weighted image using boundary-based registration. Finally, the diffusion DTI data were registered to the native structural space and masked to the appropriate size. In an additional step, FSL’s Bedpostx^68,69^ was applied to estimate the uncertainty of each fiber in each voxel for three possible fiber orientations, as was done for the HCP dataset. The transformation between different spaces was conducted in the same way as for the HCP dataset.

### Construction of tract-geometry coupling (TGC)

#### Cortical geometric eigenmodes

We followed the procedure from a previous study to obtain the cortical geometric eigenmodes ^24^. The spatial aspect of brain structure satisfies the Laplacian eigenvalue problem, also known as the Helmholtz equation, defined in the following equation,

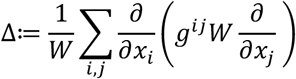

where *x_i_*, *x_i_* are the local coordinates, *g^ij^* is the inverse of the inner product metric tensor *g_ij_* ≔< 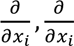>, *W* ≔ 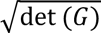, *det* denoted the determinant, and *G* ≔ (*g_ij_*).

We derived the geometric eigenmodes of the cortical surfaces by solving the eigen-decomposition problem *ΔU* = *U*Λ, where *U* is composed of eigenvectors *u*_i_ and the corresponding eigenvalue *λ_i_*. Specifically, we used a triangular surface mesh representation of the gray-white matter interface cortical surface, comprising 32,492 vertices in each hemisphere. The surface used for the HCP analysis was from the published version (https://github.com/ThomasYeoLab/CBIG/tree/master/data/templates/surface/fs_LR_32k). Note that the templates for the analysis of IMAGEN data were calculated by averaging the reconstructed surface of all subjects in FU3.

#### Connectivity blueprints

To calculate the connectivity blueprints for each subject, we performed probabilistic tractography using FSL’s probtrackx2 ^68,69^ accelerated by using GPUs ^70^. Specifically, the white surface was set as a seed region tracking to the rest of the brain with the ventricles removed and down-sampled to 3 mm resolution. The pial surface was used as a stop mask to prevent streamlined from crossing sulci. Each vertex was sampled 5000 times (5000 trackings) based on the orientation probability model for each voxel, with a curvature threshold of 0.2, a step length of 0.5 mm, and a number of steps of 3200. This resulted in a (*whole-surface vertices*) × (*whole-brain voxels*) matrix.

We utilized probabilistic tractography to quantify the tract-to-cortex patterns, i.e., the tract reachability for each tract segmented by TractSeg ^25^. This approach accounted for the inherent uncertainty in measuring and mapping white matter tracts, providing a more comprehensive and accurate representation of the brain’s structural connectivity. Specifically, we reconstructed 72 fiber bundles using a pre-trained deep-learning model, TractSeg ^25^, and down-sampled the resulting tract masks to 3 mm resolution, yielding a (*tracts*) × (*whole-brain voxels*) matrix. Then the connectivity blueprints were generated by the product of this tract matrix, and the whole-brain connectivity matrix was formed and normalized ^26^. The columns of the resulting (*tracts*) × (*whole-surface vertices*) matrix showed the connectivity distribution pattern of each cortical vertex, while the rows revealed the cortical projection patterns of the tracts, i.e., the tract reachability maps. Note that the superior cerebellar peduncle (SCP), middle cerebellar peduncle (MCP), inferior cerebellar peduncle (ICP), and fornix (FX) were disregarded in the follow-up analysis due to the absence of cortical projections, and the overall corpus callosum (CC – all) was disregarded because its 7 subsections were available.

#### Construction of tract-geometry coupling (TGC)

We used the geometric eigenmodes to decompose the reachability maps for the tracts. The orthogonal eigenmodes form a complete basis set, with the corresponding eigenvalues ordered sequentially according to the spatial frequency of the spatial patterns of each mode. The reachability maps of the tracts for each subject were reconstructed by the weighted sum of the geometric eigenmodes,

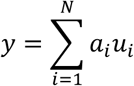

where *a_i_* is the amplitude of eigenmode *i* in explaining the data, *u_i_* is the *i*^th^ eigenmode, and *N* is the number of eigenmodes used. The tract-geometry coupling (TGC) was defined by *TGC* ≔<*a*_1_, *a*_2_, …, *a_N_* >. We used *N* = 200 in the Helmholtz equation in the current analyses. Then, we used the obtained amplitudes to calculate the reconstructed tract reachability maps. We quantified the accuracy of the reconstruction by calculating the correlation between the empirical and reconstructed data.

#### Specificity of TGC

To test the specificity of the TGC, we used two approaches of spin test to randomize geometric eigenmodes and reconstruct tract reachability. We shuffled the eigenmodes to generate 1,000 surrogate maps using BrainSMASH ^29^ and eigenstrapping ^30^ and compared the reconstruction accuracy between those using actual geometric eigenmodes.

### Characterization of TGC

#### Individual identification

To explore whether the TGC is generally stable across the subjects, we calculated the correlation coefficient of TGC between the two scans of the same subject in the HCP, a cohort that underwent test and retest scanning over several days (*n* = 44, 12 males; mean age, 30.35±3.76; age range, 22-35).

Moreover, we wanted to investigate whether it could capture individual differences between the subjects ^31^. First, a database was created that consisted of all the individual subjects’ TGCs from the second scanning, 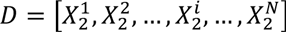, where 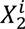 is the TGC of the subject *i* and the *N* denotes the number of subjects. To identify the current target matrix 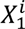, the similarity between all matrices in *D* was computed, and the predicted identity was the one with the maximum similarity score. Similarity was defined as the Pearson correlation between the target matrix and each of the database matrices. The TGC for each tract was repeated to conduct the same procedure for each tract separately.

#### Heritability of the TGC

We evaluated the heritability of the TGC using APACE ^71^. We selected 194 twins from the HCP database (102 monozygotic and 92 dizygotic pairs; mean age, 29.96±2.96; age range, 22-35). The heritability of each eigenmode was obtained by averaging the corresponding values. The difference in heritability between the low-frequency and high-frequency eigenmodes of the association, commissure, and projection tracts was also assessed.

#### Behavioral data of the HCP

We tested whether the TGC could predict behavior at the individual level. 58 behavioral scores from the HCP were considered. Because many behavioral scores were correlated, we performed a principal component analysis that was consistent with a previous study ^32^, deriving components explaining differing aspects of behavior. The top three components explaining the greatest variance were retained and interpreted as being related to (1) cognition, (2) life dissatisfaction and (3) emotional recognition.

#### Behavioral data of IMAGEN

We included 4 measures of dysfunctional behaviors in our analyses, selected from all available mental health-relevant assessments taken from participants in the IMAGEN study (follow-up3). This consisted of 2 measures of psychiatric symptoms (Community Assessment of Psychic Experiences [CAPE], Strengths and Difficulties Questionnaires [SDQ]) and 2 measures of addictive behavior from the Fagerstrom Test for Nicotine Dependence (FTND) and the Alcohol Use Disorders Identification Test (AUDIT). The primary questions of interest were regarding lifetime alcohol use and lifetime cigarette consumption.

CAPE is a self-report tool which can be used to measure the subclinical psychosis phenotype with good reliability and validity ^72^. It can capture not only positive psychotic experiences but also attenuated negative symptoms. Here we used the Total Score CAPE, which is the sum of Positive Dimension Frequency Score, Positive Dimension Distress Score, Depressive Dimension Frequency Score, Depressive Dimension Distress Score, Negative Dimension Frequency Score, and Negative Dimension Distress Score. The SDQ is a reliable and valid measure of youth emotional and behavioral symptoms. It can be used to assess 5 dimensions of youth pro-social and antisocial behaviors: emotional symptoms, conduct problems, hyperactivity/inattention, peer relationship problems, and prosocial behavior ^73^. Here we used sebdtot, which is the total difficulties score.

For the addictive behavior, we used the FTND and the AUDIT score. The FTND is used to assess nicotine dependence and smoking frequency ^74^. Here we used ftnd_sum, which is the sum of ftnd1 to ftnd6. Higher scores indicate that withdrawal symptoms from quitting tobacco are likely to be stronger. The AUDIT was developed and validated by the World Health Organization to assist with the brief assessment of alcohol use disorders and was specifically designed for international use. Here we used the sum of the Frequency and Hazardous Alcohol Use, Dependence Symptoms and Harmful Alcohol Use.

In addition, we added another six psychopathology symptoms in the IMAGEN dataset, which describe depression (depband), specific fear (spphband), generalised anxiety (dcgena), obessive compulsive disorder (ocdband), eating disorder (eatband), and exceptionally stressful event (ptsdband).

#### Behavior prediction models

We utilized kernel ridge regression (KRR) to predict the 3 behavioral components from the HCP data and other behavioral measures from the IMAGEN dataset, which has shown strong behavioral prediction performance ^32^. An L2-regularization term was used to reduce overfitting.

The TGC was used as the independent feature in the regression model. We performed 10-fold nested cross-validation for each behavior measure. For each test fold, the KRR parameters were estimated from the remain nine training folds and the best L2-regularization parameter was selected. Sixty random replications of 10-fold nested cross-validation were performed to mitigate the sensitivity led by a single cross-validation.

Age and sex were regressed from the behavioral measures. Regression was performed on the training folds, and regression coefficients were applied to the test fold. The accuracy of each model was defined as the correlation between the predicted scores of the test participants and their actual scores within each test fold. These correlations were then averaged across 10 test folds and replications for each behavioral measure. 1,000 iterations of permutation test were performed to assess the significance.

In addition, tract reachability was also used to predict behavior and its performance was compared to those using TGC.

#### Model interpretation

For each component score, we interpreted the importance of features by using the Haufe transformation ^33^. In brief, for the given prediction model, the covariance was calculated between each feature and the predicted behavior score for each subject. Positive feature importance scores indicated that greater brain feature values were associated with greater predicted behavior values.

The transformation yielded a 7,200-length vector, which was rearranged back into a 200 × 36 matrix. We also calculated the norm across the eigenmodes or tracts to derive the importance values of eigenmodes and tracts.

### High-low frequency ratio of TGC

#### High-low frequency ratio

Following Preti et al. ^34^, we implemented graph signal filtering to decompose the tract reachability maps into one part that was well coupled with geometry (i.e., represented by low-frequency eigenmodes) and one that was less coupled (i.e., high-frequency eigenmodes). The cut-off frequency *C* was selected after splitting the spectrum into two portions with equal energy (median-split) based on the average energy spectral density. The matrix *U^low^* contains the first *C* eigenmodes (columns of *U*) complemented with *N* − *C* zero columns. In contrast, the matrix *U^high^* contains the first *C* zero columns followed by the *N* − *C* last eigenmodes. Therefore, the filtered patterns were obtained using the following equations:

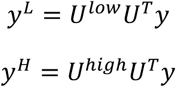

As a measure of the tract-geometry-coupling of a specific tract, we defined the high-low frequency ratio, i.e., the ratio between the norm of *y^H^* and *y^L^* across the vertices where the white matter tract reached the surface. A high-low frequency ratio map was obtained for each tract as the ratio between *y^H^* and *y^L^*. The average map was generated by averaging the ratio maps across tracts.

#### Functional decoding of the high-low frequency ratio map

We projected the meta-analytical task-based activation along the average high-low frequency ratio map to conduct functional decoding. Here, we chose 590 terms related to specific cognitive processes that had been selected in previous publications ^75^. The activation maps for the 590 cognitive terms were downloaded from the NeuroSynth database ^39^ (https://neurosynth.org/). We generated 20 binarized masks at five-percentile increments of the high-low frequency ratio map. Each of these 20 maps, ranging from 0-5% to 95%-100%, were used as inputs to the subsequent meta-analysis. Each function term had a mean activation z-score per bin. The terms included in the visualization were those that had the highest five z-scores in each bin. A significance threshold of *z* > 0.5 was added as a visualization constraint.

### Functional associations of TGC

#### HCP Task activation data

We used the task fMRI data that were projected into 2 mm standard CIFTI grayordinates space. The task fMRI contained 86 contrasts from seven task domains ^76^, labelled as EMOTION, GAMBLING, LANGUAGE, MOTOR, RELATIONAL, SOCIAL, and WM. Paired negative contrasts are redundant for the regression modelling and were thus excluded ^77^, resulting in 47 contrasts for further analysis. The z-statistic for each contrast was used to assess the functional activations.

#### Correlate activation map with TGC

We explored the relationship between task activation and TGC by first correlating each TGC with each cortical vertex. A significant association was determined with a *p* < 0.05 after correcting for multiple comparisons using FDR, and the number of significant vertices for each TGC was counted to represent its association with the activation map. The similarity between activation maps was measured using the resultant TGCs. We also computed the correlation between the above-mentioned uncorrected correlation matrix and the activation map. TGCs that had a greater correlation value indicated that the TGCs had a stronger association with the activation map.

## Data availability

HCP data are available at https://db.humanconnectome.org/. IMAGEN data are available at https://imagen2.cea.fr. The Yeo 7-network atlas can be downloaded at https://github.com/ThomasYeoLab/CBIG/tree/master/stable_projects/brain_parcellation/Yeo2011_fcMRI_clustering/1000subjects_reference/Yeo_JNeurophysiol11_SplitLabels/fs_LR32k. The activated maps of the cognitive terms can be downloaded from the NeuroSynth database (https://neurosynth.org/).

## Code availability

The HCP-Pipeline can be found at https://github.com/Washington-University/HCPpipelines. The neuroimaging preprocessing software used for the other datasets is freely available (FreeSurfer v7.3.2, http://surfer.nmr.mgh.harvard.edu/ and FSL v6.0.7, https://fsl.fmrib.ox.ac.uk/fsl/fslwiki). The brain maps were presented using BrainSpace (https://brainspace.readthedocs.io/) and Workbench v1.5.0 (https://www.humanconnectome.org/software/connectome-workbench/). The code to generate the results in this work is available at https://github.com/FANLabCASIA/D_TGC_tract-geometry-coupling.git.

## Acknowledgments

This work was partially supported by STI2030-Major Projects (Grant No. 2021ZD0200200), the Natural Science Foundation of China (Grant Nos. 82072099, 82202253, 62250058), and the China Postdoctoral Science Foundation (2022M722915). Data were provided in part by the Human Connectome Project, WU-Minn Consortium (Principal Investigators: David Van Essen and Kamil Ugurbil; 1U54MH091657), which was funded by the 16 NIH Institutes and Centers that support the NIH Blueprint for Neuroscience Research and by the McDonnell Center for Systems Neuroscience at Washington University; the European Union-funded FP6 Integrated Project IMAGEN (Reinforcement-related behaviour in normal brain function and psychopathology) (LSHM-CT-2007-037286), the Horizon 2020 funded ERC Advanced Grant ‘STRATIFY’ (Brain network based stratification of reinforcement-related disorders) (695313), Horizon Europe ‘environMENTAL’, grant no: 101057429, UK Research and Innovation (UKRI) Horizon Europe funding guarantee (10041392 and 10038599), Human Brain Project (HBP SGA 2, 785907, and HBP SGA 3, 945539), the Chinese government via the Ministry of Science and Technology (MOST). The German Center for Mental Health (DZPG), the Bundesministerium für Bildung und Forschung (BMBF grants 01GS08152; 01EV0711; Forschungsnetz AERIAL 01EE1406A, 01EE1406B; Forschungsnetz IMAC-Mind 01GL1745B), the Deutsche Forschungsgemeinschaft (DFG grants SM 80/7-2, SFB 940, TRR 265, NE 1383/14-1, 186318919 [FOR 1617], 178833530 [SFB 940], 386691645 [NE 1383/14-1], 402170461 [TRR 265], 454245598 [IRTG 2773]), the Medical Research Foundation and Medical Research Council (grants MR/R00465X/1 and MR/S020306/1), the National Institutes of Health (NIH) funded ENIGMA-grants 5U54EB020403-05, 1R56AG058854-01 and U54 EB020403 as well as NIH R01DA049238, the National Institutes of Health, Science Foundation Ireland (16/ERCD/3797). NSFC grant 82150710554. Further support was provided by grants from: the ANR (ANR-12-SAMA-0004, AAPG2019 - GeBra), the Eranet Neuron (AF12-NEUR0008-01 WM2NA; and ANR-18-NEUR00002-01 - ADORe), the Fondation de France (00081242), the Fondation pour la Recherche Médicale (DPA20140629802), the Mission Interministérielle de Lutte-contre-les-Drogues-et-les-Conduites-Addictives (MILDECA), the Assistance-Publique-Hôpitaux-de-Paris and INSERM (interface grant), Paris Sud University IDEX 2012, the Fondation de l’Avenir (grant AP-RM-17-013), the Fédération pour la Recherche sur le Cerveau, the Ile de France region (grant QIM VEAVE conventtion 23002745-23002747). The authors appreciate the English language and editing assistance of Rhoda E. and Edmund F. Perozzi, PhDs.

