## Supplemental Table 1 for "Mapping the coupling between tract reachability and cortical geometry of the human brain"

**Left hemisphere**

| Behavior name | r | p | Behavior description | Dataset |
|---|---|---|---|---|
| PicSeq_Unadj | 0.1246 | 1.79E-04 | Episodic Memory (Picture Sequence Memory) | HCP |
| CardSort_Unadj | 0.2429 | 1.50E-13 | Executive Function/Cognitive Flexibility (Dimensional Change Card Sort) | HCP |
| Flanker_Unadj | 0.1226 | 2.26E-04 | Executive Function/Inhibition (Flanker Task) | HCP |
| PMAT24_A_CR | 0.2607 | 1.88E-15 | Fluid Intelligence (Penn Progressive Matrices) | HCP |
| ReadEng_Unadj | 0.2358 | 7.77E-13 | Language/Reading Decoding (Oral Reading Recognition) | HCP |
| PicVocab_Unadj | 0.263 | 1.04E-15 | Language/Vocabulary Comprehension (Picture Vocabulary) | HCP |
| ProcSpeed_Unadj | 0.1128 | 7.01E-04 | Processing Speed (Pattern Completion Processing Speed) | HCP |
| DDisc_AUC_40K | 0.2315 | 2.03E-12 | Self-regulation/Impulsivity (Delay Discounting) | HCP |
| VSPLOT_TC | 0.248 | 4.40E-14 | Spatial Orientation (Variable Short Penn Line Orientation Test) | HCP |
| SCPT_SEN | 0.0911 | 6.23E-03 | Sustained Attention (Short Penn Continuous Performance Test) | HCP |
| SCPT_SPEC | 0.0848 | 1.09E-02 | Sustained Attention (Short Penn Continuous Performance Test) | HCP |
| IWRD_TOT | 0.1125 | 7.22E-04 | Verbal Episodic Memory (Penn Word Memory Test) | HCP |
| ListSort_Unadj | 0.1608 | 1.24E-06 | Working Memory (List Sorting) | HCP |
| MMSE_Score | 0.0566 | 8.96E-02 | Cognitive Status (Mini Mental Status Exam) | HCP |
| PSQI_Score | 0.0783 | 1.88E-02 | Sleep (Pittsburgh Sleep Questionnaire) | HCP |
| Endurance_Unadj | 0.274 | 5.73E-17 | Endurance (2 minute walk test) | HCP |
| GaitSpeed_Comp | -0.001 | 9.66E-01 | Locomotion (4-meter walk test) | HCP |
| Dexterity_Unadj | 0.2604 | 2.03E-15 | Dexterity (9-hole Pegboard) | HCP |
| Strength_Unadj | 0.5763 | 9.33E-81 | Strength (Grip Strength Dynamometry) | HCP |

| | | | | |
|---|---|---|---|---|
| Odor_Unadj | 0.0591 | 7.65E-02 | Olfaction (Odor Identification Test) | HCP |
| PainInterf_Tscore | 0.0482 | 1.49E-01 | Pain (Pain Intensity and Interference Surveys) | HCP |
| Taste_Unadj | 0.1759 | 1.10E-07 | Taste (Taste Intensity Test) | HCP |
| Mars_Final | 0.0032 | 9.25E-01 | Contrast Sensitivity (Mars Contrast Sensitivity) | HCP |
| Emotion_Task_Face_Acc | 0.0797 | 1.68E-02 | Emotion | HCP |
| Language_Task_Math_Avg_Difficulty_Level | 0.1519 | 4.73E-06 | Language | HCP |
| Language_Task_Story_Avg_Difficulty_Level | 0.1816 | 4.13E-08 | Language | HCP |
| Relational_Task_Acc | 0.2293 | 3.37E-12 | Relational | HCP |
| Social_Task_Perc_Random | 0.0715 | 3.19E-02 | Social | HCP |
| Social_Task_Perc_TOM | 0.0724 | 3.00E-02 | Social | HCP |
| WM_Task_Acc | 0.2151 | 7.00E-11 | Working Memory | HCP |
| NEOFAC_A | 0.179 | 6.48E-08 | Five Factor Model (NEO-FFI) Factor Summary Scores | HCP |
| NEOFAC_O | 0.0647 | 5.23E-02 | Five Factor Model (NEO-FFI) Factor Summary Scores | HCP |
| NEOFAC_C | 0.1402 | 2.43E-05 | Five Factor Model (NEO-FFI) Factor Summary Scores | HCP |
| NEOFAC_N | 0.0672 | 4.38E-02 | Five Factor Model (NEO-FFI) Factor Summary Scores | HCP |
| NEOFAC_E | 0.0555 | 9.62E-02 | Five Factor Model (NEO-FFI) Factor Summary Scores | HCP |
| ER40_CR | 0.1064 | 1.39E-03 | Emotion Recognition (Penn Emotion Recognition Test) | HCP |

| | | | | |
|---|---|---|---|---|
| ER40ANG | 0.0725 | 2.96E-02 | Emotion Recognition (Penn Emotion Recognition Test) | HCP |
| ER40FEAR | 0.1144 | 5.86E-04 | Emotion Recognition (Penn Emotion Recognition Test) | HCP |
| ER40HAP | 0.049 | 1.42E-01 | Emotion Recognition (Penn Emotion Recognition Test) | HCP |
| ER40NOE | 0.0147 | 6.59E-01 | Emotion Recognition (Penn Emotion Recognition Test) | HCP |
| ER40SAD | 0.1448 | 1.30E-05 | Emotion Recognition (Penn Emotion Recognition Test) | HCP |
| AngAffect_Unadj | 0.0047 | 8.87E-01 | Negative Affect (Sadness, Fear, Anger) | HCP |
| AngHostil_Unadj | 0.0645 | 5.29E-02 | Negative Affect (Sadness, Fear, Anger) | HCP |
| AngAggr_Unadj | 0.2209 | 2.08E-11 | Negative Affect (Sadness, Fear, Anger) | HCP |
| FearAffect_Unadj | 0.051 | 1.26E-01 | Negative Affect (Sadness, Fear, Anger) | HCP |
| FearSomat_Unadj | 0.0704 | 3.48E-02 | Negative Affect (Sadness, Fear, Anger) | HCP |
| Sadness_Unadj | 0.0603 | 7.05E-02 | Negative Affect (Sadness, Fear, Anger) | HCP |
| LifeSatisf_Unadj | 0.1652 | 6.21E-07 | Psychological Well-being (Positive Affect, Life Satisfaction, Meaning and Purpose) | HCP |
| MeanPurp_Unadj | 0.0587 | 7.85E-02 | Psychological Well-being (Positive Affect, Life Satisfaction, Meaning and Purpose) | HCP |
| PosAffect_Unadj | 0.1206 | 2.89E-04 | Psychological Well-being (Positive Affect, Life Satisfaction, Meaning and Purpose) | HCP |
| Friendship_Unadj | 0.0359 | 2.81E-01 | Social Relationships (Social Support, Companionship, Social Distress, Positive Social Development) | HCP |
| Loneliness_Unadj | 0.0362 | 2.79E-01 | Social Relationships (Social Support, Companionship, Social Distress, Positive Social Development) | HCP |
| PercHostil_Unadj | 0.133 | 6.25E-05 | Social Relationships (Social Support, Companionship, Social Distress, Positive Social Development) | HCP |

| | | | | |
|---|---|---|---|---|
| PercReject_Unadj | 0.0868 | 9.14E-03 | Social Relationships (Social Support, Companionship, Social Distress, Positive Social Development) | HCP |
| EmotSupp_Unadj | 0.0865 | 9.46E-03 | Social Relationships (Social Support, Companionship, Social Distress, Positive Social Development) | HCP |
| InstruSupp_Unadj | 0.0627 | 6.01E-02 | Social Relationships (Social Support, Companionship, Social Distress, Positive Social Development) | HCP |
| PercStress_Unadj | 0.1107 | 8.76E-04 | Stress and Self Efficacy (Perceived Stress, Self-Efficacy) | HCP |
| SelfEff_Unadj | 0.1592 | 1.59E-06 | Stress and Self Efficacy (Perceived Stress, Self-Efficacy) | HCP |
| Dissatification | 0.1124 | 0.0007303 | PCA components | HCP |
| Cognation | 0.3622 | 2.72E-29 | PCA components | HCP |
| Emotion | 0.1761 | 1.056E-07 | PCA components | HCP |
| CAPE | 0.1973 | 1.115E-06 | | IMAGEN FU3 |
| ESPAD | 0.171 | 6.523E-05 | | IMAGEN FU3 |
| SDQ | 0.1979 | 1.025E-06 | | IMAGEN FU3 |
| AUDIT | 0.2002 | 7.613E-07 | | IMAGEN FU3 |
| Dissatification | 0.1086 | 0.00118 | PCA components | HCP individual version |
| Cognation | 0.3921 | 4.369E-34 | PCA components | HCP individual version |
| Emotion | 0.1086 | 0.001181 | PCA components | HCP individual version |

**Right hemisphere**

| Behavior name | r | p | Behavior description | Dataset |
|---|---|---|---|---|
| PicSeq_Unadj | 0.1593 | 1.554E-06 | Episodic Memory (Picture Sequence Memory) | HCP |
| CardSort_Unadj | 0.1731 | 1.731E-07 | Executive Function/Cognitive Flexibility (Dimer | HCP |
| Flanker_Unadj | 0.1139 | 0.0006175 | Executive Function/Inhibition (Flanker Task) | HCP |
| PMAT24_A_CR | 0.2497 | 2.903E-14 | Fluid Intelligence (Penn Progressive Matrices) | HCP |
| ReadEng_Unadj | 0.2416 | 2.012E-13 | Language/Reading Decoding (Oral Reading Rec | HCP |
| PicVocab_Unadj | 0.2418 | 1.921E-13 | Language/Vocabulary Comprehension (Picture | HCP |
| ProcSpeed_Unadj | 0.0926 | 0.0054443 | Processing Speed (Pattern Completion Processin | HCP |
| DDisc_AUC_40K | 0.1313 | 7.839E-05 | Self-regulation/Impulsivity (Delay Discounting) | HCP |
| VSPLOT_TC | 0.2655 | 5.532E-16 | Spatial Orientation (Variable Short Penn Line O | HCP |
| SCPT_SEN | 0.0304 | 0.3618559 | Sustained Attention (Short Penn Continuous Per | HCP |
| SCPT_SPEC | 0.0715 | 0.0320134 | Sustained Attention (Short Penn Continuous Per | HCP |
| IWRD_TOT | 0.1212 | 0.0002689 | Verbal Episodic Memory (Penn Word Memory | HCP |
| ListSort_Unadj | 0.1688 | 3.511E-07 | Working Memory (List Sorting) | HCP |
| MMSE_Score | 0.0692 | 0.0378195 | Cognitive Status (Mini Mental Status Exam) | HCP |
| PSQI_Score | 0.0932 | 0.0051243 | Sleep (Pittsburgh Sleep Questionnaire) | HCP |
| Endurance_Unadj | 0.2925 | 3.222E-19 | Endurance (2 minute walk test) | HCP |
| GaitSpeed_Comp | 0.0446 | 0.1814362 | Locomotion (4-meter walk test) | HCP |
| Dexterity_Unadj | 0.1965 | 2.772E-09 | Dexterity (9-hole Pegboard) | HCP |
| Strength_Unadj | 0.5716 | 3.363E-79 | Strength (Grip Strength Dynamometry) | HCP |
| Odor_Unadj | 0.0444 | 0.1833774 | Olfaction (Odor Identification Test) | HCP |
| PainInterf_Tscore | 0.068 | 0.0414972 | Pain (Pain Intensity and Interference Surveys) | HCP |
| Taste_Unadj | 0.176 | 1.072E-07 | Taste (Taste Intensity Test) | HCP |
| Mars_Final | 0.003 | 0.9288551 | Contrast Sensitivity (Mars Contrast Sensitivity) | HCP |
| Emotion_Task_Face_Acc | 0.0603 | 0.0706827 | Emotion | HCP |
| Language_Task_Math_Avg | 0.1569 | 2.235E-06 | Language | HCP |
| Language_Task_Story_Av | 0.2092 | 2.334E-10 | Language | HCP |
| Relational_Task_Acc | 0.2623 | 1.261E-15 | Relational | HCP |
| Social_Task_Perc_Random | 0.0792 | 0.0174269 | Social | HCP |
| Social_Task_Perc_TOM | 0.0735 | 0.0275597 | Social | HCP |
| WM_Task_Acc | 0.2555 | 6.999E-15 | Working Memory | HCP |
| NEOFAC_A | 0.1961 | 3.004E-09 | Five Factor Model (NEO-FFI) Factor Summary | HCP |
| NEOFAC_O | 0.1852 | 2.196E-08 | Five Factor Model (NEO-FFI) Factor Summary | HCP |
| NEOFAC_C | 0.1671 | 4.6E-07 | Five Factor Model (NEO-FFI) Factor Summary | HCP |
| NEOFAC_N | 0.1496 | 6.527E-06 | Five Factor Model (NEO-FFI) Factor Summary | HCP |

| | | | | |
|---|---|---|---|---|
| NEOFAC_E | 0.096 | 0.0039407 | Five Factor Model (NEO-FFI) Factor Summary | HCP |
| ER40_CR | 0.1155 | 0.0005162 | Emotion Recognition (Penn Emotion Recognitic | HCP |
| ER40ANG | 0.0819 | 0.0139988 | Emotion Recognition (Penn Emotion Recognitic | HCP |
| ER40FEAR | 0.0361 | 0.2790075 | Emotion Recognition (Penn Emotion Recognitic | HCP |
| ER40HAP | 0.0362 | 0.2778969 | Emotion Recognition (Penn Emotion Recognitic | HCP |
| ER40NOE | 0.0657 | 0.0487164 | Emotion Recognition (Penn Emotion Recognitic | HCP |
| ER40SAD | 0.0706 | 0.034082 | Emotion Recognition (Penn Emotion Recognitic | HCP |
| AngAffect_Unadj | 0.0474 | 0.1554327 | Negative Affect (Sadness, Fear, Anger) | HCP |
| AngHostil_Unadj | 0.028 | 0.4006647 | Negative Affect (Sadness, Fear, Anger) | HCP |
| AngAggr_Unadj | 0.1851 | 2.231E-08 | Negative Affect (Sadness, Fear, Anger) | HCP |
| FearAffect_Unadj | 0.0581 | 0.0813392 | Negative Affect (Sadness, Fear, Anger) | HCP |
| FearSomat_Unadj | 0.078 | 0.0193313 | Negative Affect (Sadness, Fear, Anger) | HCP |
| Sadness_Unadj | 0.0798 | 0.0166905 | Negative Affect (Sadness, Fear, Anger) | HCP |
| LifeSatisf_Unadj | 0.1762 | 1.04E-07 | Psychological Well-being (Positive Affect, Life : | HCP |
| MeanPurp_Unadj | 0.1028 | 0.002023 | Psychological Well-being (Positive Affect, Life : | HCP |
| PosAffect_Unadj | 0.1021 | 0.0021737 | Psychological Well-being (Positive Affect, Life : | HCP |
| Friendship_Unadj | 0.0315 | 0.3456438 | Social Relationships (Social Support, Companioı | HCP |
| Loneliness_Unadj | 0.0246 | 0.4616369 | Social Relationships (Social Support, Companioı | HCP |
| PercHostil_Unadj | 0.1087 | 0.0010877 | Social Relationships (Social Support, Companioı | HCP |
| PercReject_Unadj | 0.0505 | 0.1299345 | Social Relationships (Social Support, Companioı | HCP |
| EmotSupp_Unadj | 0.0595 | 0.0742888 | Social Relationships (Social Support, Companioı | HCP |
| InstruSupp_Unadj | 0.0656 | 0.0491258 | Social Relationships (Social Support, Companioı | HCP |
| PercStress_Unadj | 0.1068 | 0.0013403 | Stress and Self Efficacy (Perceived Stress, Self-E | HCP |
| SelfEff_Unadj | 0.0935 | 0.004977 | Stress and Self Efficacy (Perceived Stress, Self-E | HCP |
| Dissatification | 0.091 | 0.0062857 | PCA components | HCP |
| Cognation | 0.3711 | 9.16E-31 | PCA components | HCP |
| Emotion | 0.1634 | 8.27E-07 | PCA components | HCP |
| CAPE | 0.1316 | 3.87E-03 | | IMAGEN FU3 |
| ESPAD | 0.1443 | 0.000771 | | IMAGEN FU3 |
| SDQ | 0.1536 | 0.0007327 | | IMAGEN FU3 |
| AUDIT | 0.2606 | 9.009E-11 | | IMAGEN FU3 |
