## Supplemental Table 2 for "Mapping the coupling between tract reachability and cortical geometry of the human brain"

| fiber | fiber type | fiber decoupling index | dataset |
|---|---|---|---|
| CC_2 | Commissural | 0.306328 | HCP |
| T_PAR | Projection | 0.321475 | HCP |
| ST_OCC | Projection | 0.322565 | HCP |
| ST_PAR | Projection | 0.332469 | HCP |
| ST_PREF | Projection | 0.365339 | HCP |
| T_OCC | Projection | 0.395788 | HCP |
| IFO | Association | 0.397685 | HCP |
| CC_7 | Commissural | 0.417459 | HCP |
| OR | Projection | 0.427539 | HCP |
| ST_FO | Projection | 0.428235 | HCP |
| T_PREF | Projection | 0.431676 | HCP |
| CC_6 | Commissural | 0.433756 | HCP |
| ST_PREC | Projection | 0.45696 | HCP |
| T_PREC | Projection | 0.467145 | HCP |
| FPT | Projection | 0.486593 | HCP |
| CST | Projection | 0.503551 | HCP |
| POPT | Projection | 0.506789 | HCP |
| ILF | Association | 0.539504 | HCP |
| ST_POSTC | Projection | 0.541108 | HCP |
| CC_5 | Commissural | 0.608956 | HCP |
| AF | Association | 0.611131 | HCP |
| T_POSTC | Projection | 0.626733 | HCP |
| CC_1 | Commissural | 0.635077 | HCP |
| MLF | Association | 0.646415 | HCP |
| ST_PREM | Projection | 0.651032 | HCP |
| CC_4 | Commissural | 0.659528 | HCP |
| CG | Association | 0.669789 | HCP |
| T_PREM | Projection | 0.672586 | HCP |
| ATR | Projection | 0.673706 | HCP |
| CA | Commissural | 0.765183 | HCP |
| STR | Projection | 0.779249 | HCP |
| CC_3 | Commissural | 0.792167 | HCP |
| SLF_II | Association | 0.817246 | HCP |
| SLF_III | Association | 0.820734 | HCP |
| UF | Association | 0.869119 | HCP |
| SLF_I | Association | 0.998674 | HCP |
| **fiber** | **fiber type** | **fiber decoupling index** | |
| T_PAR | Projection | 0.315302 | IMAGEN FU3 |
| ST_PAR | Projection | 0.319779 | IMAGEN FU3 |
| ST_OCC | Projection | 0.328751 | IMAGEN FU3 |
| ST_FO | Projection | 0.375762 | IMAGEN FU3 |
| CC_2 | Commissural | 0.376059 | IMAGEN FU3 |
| T_OCC | Projection | 0.393799 | IMAGEN FU3 |
| ST_PREF | Projection | 0.414265 | IMAGEN FU3 |
| OR | Projection | 0.424568 | IMAGEN FU3 |
| CC_6 | Commissural | 0.428765 | IMAGEN FU3 |
| ST_PREC | Projection | 0.461122 | IMAGEN FU3 |
| FPT | Projection | 0.470343 | IMAGEN FU3 |
| IFO | Association | 0.470989 | IMAGEN FU3 |
| T_PREC | Projection | 0.482773 | IMAGEN FU3 |
| T_PREF | Projection | 0.490755 | IMAGEN FU3 |
| ST_POSTC | Projection | 0.515112 | IMAGEN FU3 |
| AF | Association | 0.527655 | IMAGEN FU3 |
| ILF | Association | 0.536842 | IMAGEN FU3 |
| POPT | Projection | 0.546656 | IMAGEN FU3 |

| | | | |
|---|---|---|---|
| CC_7 | Commissural | 0.547672 | IMAGEN FU3 |
| CST | Projection | 0.555256 | IMAGEN FU3 |
| MLF | Association | 0.57168 | IMAGEN FU3 |
| T_POSTC | Projection | 0.590652 | IMAGEN FU3 |
| CC_1 | Commissural | 0.659862 | IMAGEN FU3 |
| CC_4 | Commissural | 0.679527 | IMAGEN FU3 |
| CC_5 | Commissural | 0.688909 | IMAGEN FU3 |
| ST_PREM | Projection | 0.69005 | IMAGEN FU3 |
| T_PREM | Projection | 0.715173 | IMAGEN FU3 |
| ATR | Projection | 0.752548 | IMAGEN FU3 |
| SLF_II | Association | 0.791486 | IMAGEN FU3 |
| SLF_III | Association | 0.806376 | IMAGEN FU3 |
| CG | Association | 0.815205 | IMAGEN FU3 |
| CC_3 | Commissural | 0.850979 | IMAGEN FU3 |
| CA | Commissural | 0.852757 | IMAGEN FU3 |
| STR | Projection | 0.873215 | IMAGEN FU3 |
| SLF_I | Association | 0.984462 | IMAGEN FU3 |
| UF | Association | 1.053766 | IMAGEN FU3 |
