## Supplemental Figures for "Mapping the coupling between tract reachability and cortical geometry of the human brain"

### Supplementary Figures

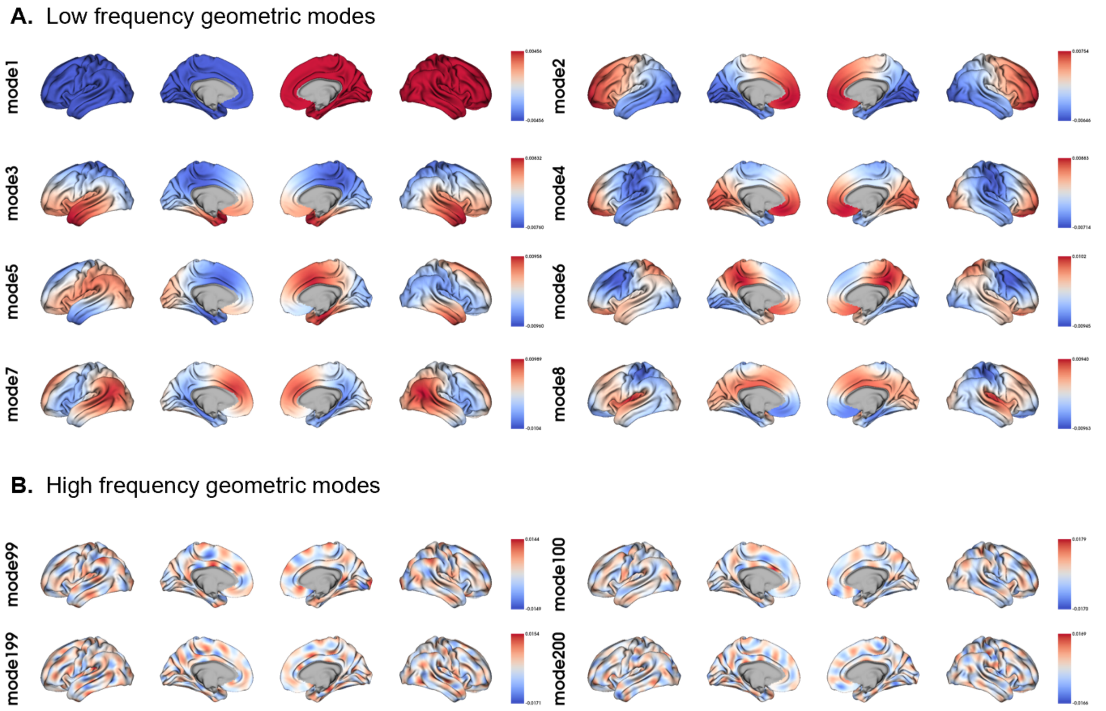


**Figure S1. Geometric eigenmodes decomposed from the human brain surface. (A)** Visualization of example low frequency geometric modes. **(B)** Visualization of example high frequency geometric modes.

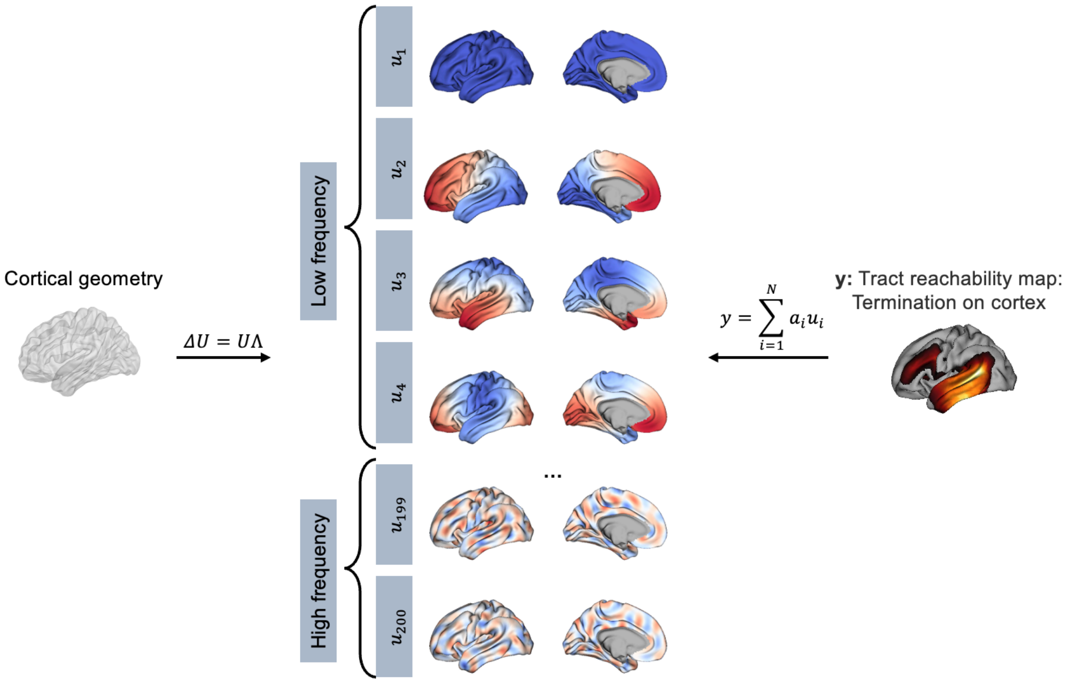


**Figure S2. The calculation of tract-geometry coupling (TGC).** A population-averaged template of the cortical surface was used to map cortical eigenmodes, which were used to reconstruct cortical projection patterns of fiber bundles. The fiber bundles were delineated with TractSeg, and their reachability to the cortex was quantified using the connectivity blueprint approach. The rows of the matrix provided the cortical projection patterns of the white matter tracts, which is explicitly referred to as the "tract reachability map". The tract reachability map of each tract was reconstructed using 200 eigenmodes for each subject using general linear models (GLM).

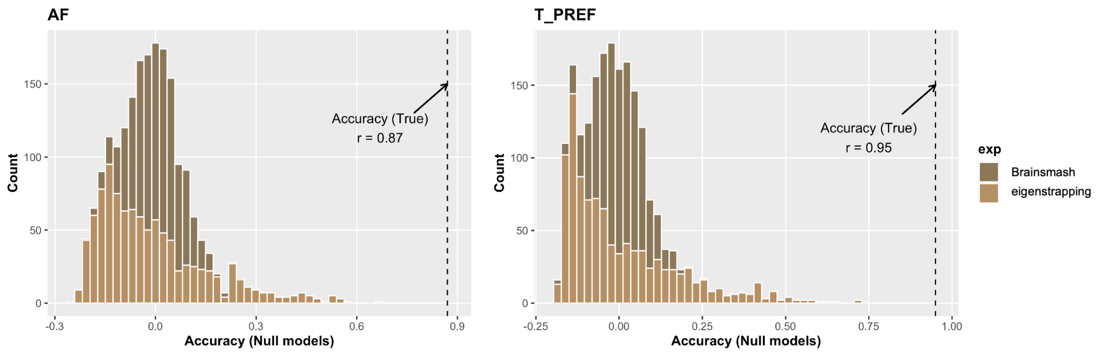


**Figure S3. Specificity of geometric eigenmodes.** We used two approaches to randomize geometric eigenmodes and reconstruct tract reachability. We found the accuracy was significantly lower than actual eigenmodes. Here we showed results of two example white matter tracts, i.e., arcuate fasciculus (AF) and thalamo-prefrontal tract (T_PREF).

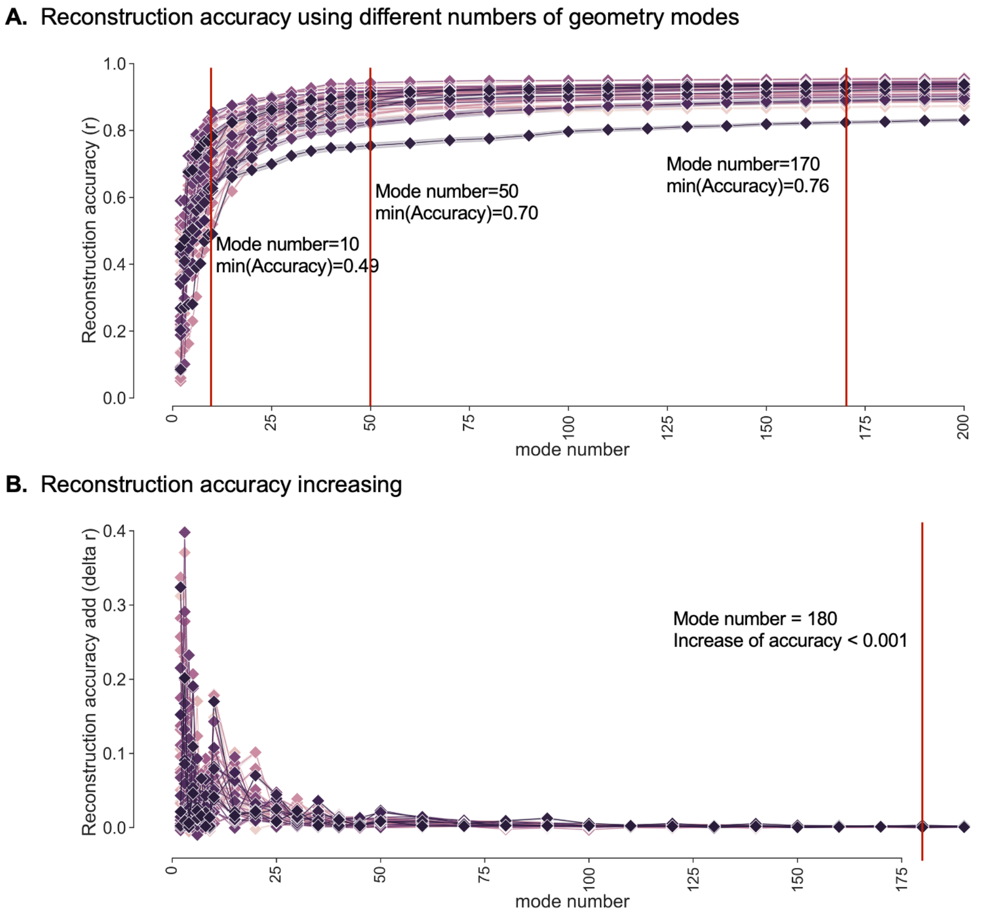


**Figure S4. Reconstruction accuracy using different numbers of geometric eigenmodes. (A)** We found that reconstruction accuracy increased with an increasing number of eigenmodes across all tract reachability maps, with $r > 0.5$ already achieved using just $N = 10$ modes and $r > 0.7$ using $N = 50$. **(B)** The increasement of reconstructed accuracy is less than 0.001 when using more than 180 geometric eigenmodes.

**A.** Reconstruction accuracy using individual eigenmodes

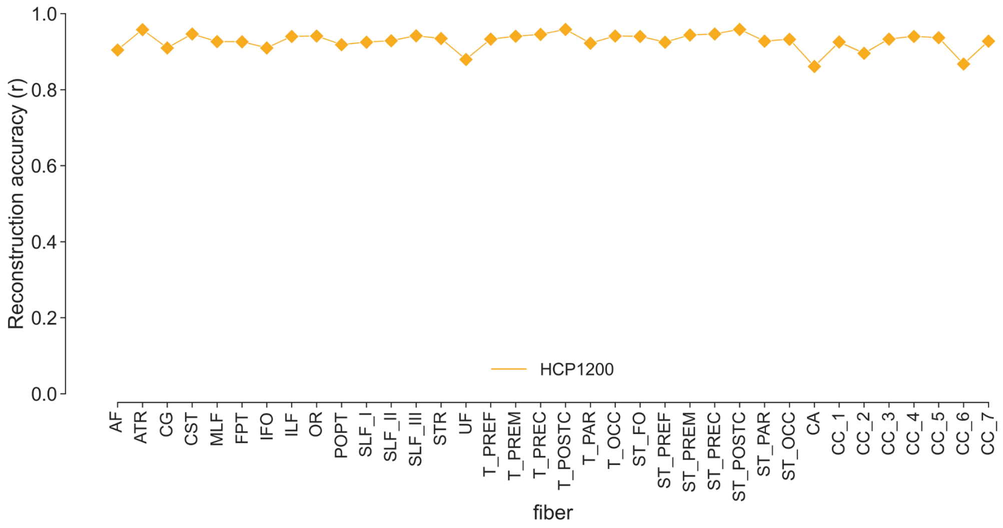


**B.** Correlation between TGC (group eigenmodes) and TGC (individual eigenmodes)

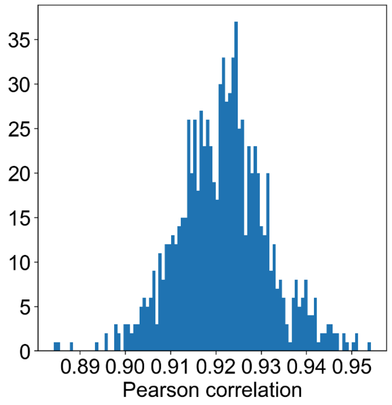


**Figure S5. Results using individual geometric eigenmodes. (A)** Reconstruction of tract reachability using individual eigenmodes remained very high. **(B)** The high similarity between TGC using group eigenmodes and individual eigenmodes.

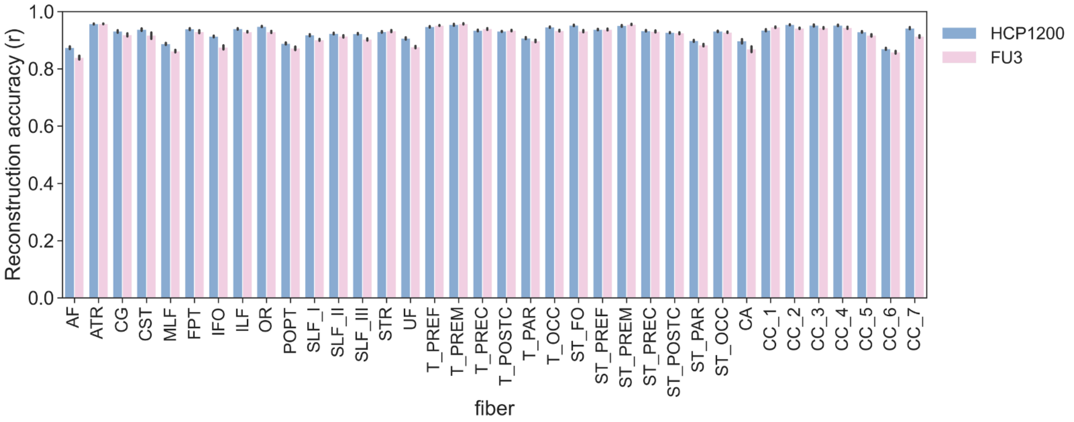


**Figure S6. Reconstruction of tract reachability maps of right hemisphere.** Reconstruction accuracy of tracts in the HCP and IMAGEN datasets. High accuracy (all correlation coefficients > 0.8) were observed in both datasets.

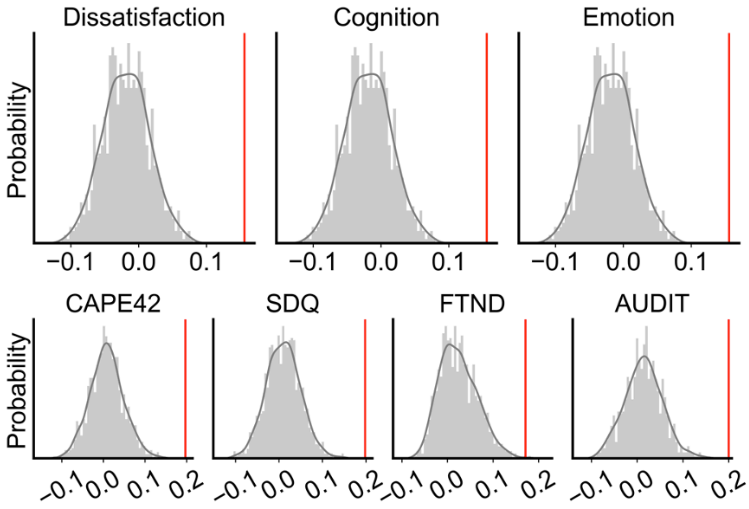


**Figure S7. The permutation test of behavior prediction in HCP and IMAGEN dataset.**

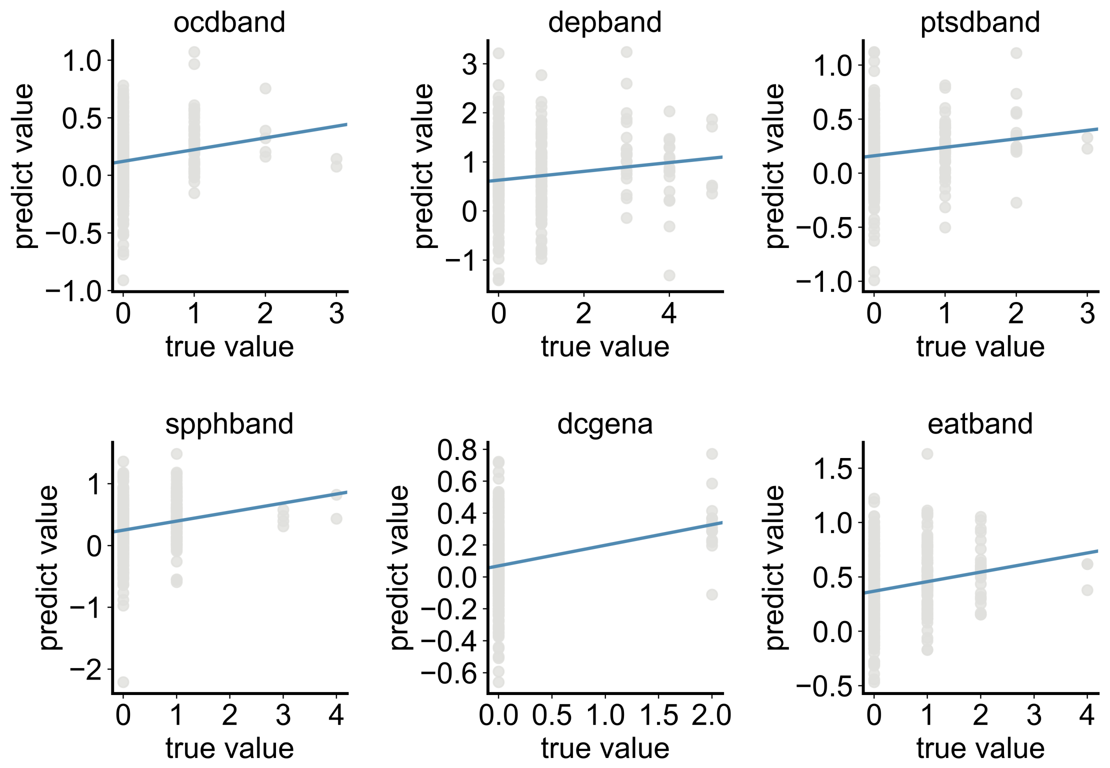


**Figure S8. TGC could predict psychopathology symptoms in the IMAGEN dataset.** depband: $r = 0.13$, $p_{\text{permutation}} = 0.25$; spphband: $r = 0.22$, $p_{\text{permutation}} = 0$; dcgena: $r = 0.23$, $p_{\text{permutation}} = 0.002$; ocdband: $r = 0.19$, $p_{\text{permutation}} = 0.006$; eatband: $r = 0.19$, $p_{\text{permutation}} = 0$; ptsdband: $r = 0.15$, $p_{\text{permutation}} = 0.014$.

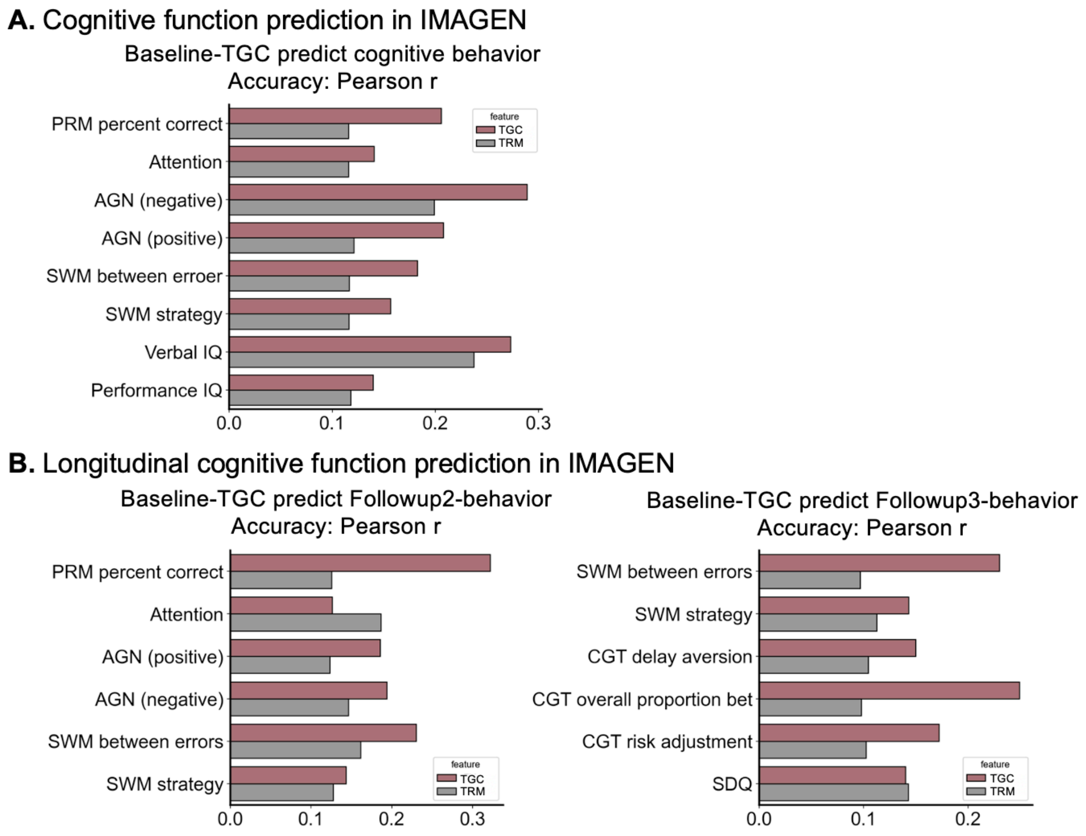


**Figure S9. Behavior prediction accuracy of TGC and tract reachability.** The tract reachability maps (TRM), i.e., the reachability map for all white matter tracts was used to predict behaviors. **(A)** TGC and TRM both could predict cognitive behaviors in the IMAGEN dataset and TGC has a better performance (PRM percent correct: $r = 0.21$, $p = 0.001$ [TGC]; $r = 0.12$, $p = 0.006$ [TRM]; Attention: $r = 0.14$, $p = 0.001$ [TGC]; $r = 0.12$, $p = 0.015$ [TRM]; AGN negative: $r = 0.29$, $p = 0.001$ [TGC]; $r = 0.20$, $p = 0.001$ [TRM]; AGN positive: $r = 0.21$, $p = 0.001$ [TGC]; $r = 0.12$, $p = 0.006$ [TRM]; SWM between errors: $r = 0.18$, $p = 0.001$ [TGC]; $r = 0.12$, $p = 0.065$ [TRM]; SWM strategy: $r = 0.16$, $p = 0.001$ [TGC]; $r = 0.12$, $p = 0.024$ [TRM]; Verbal ID: $r = 0.27$, $p = 0.001$ [TGC]; $r = 0.24$, $p = 0.001$ [TRM]; Performance IQ: $r = 0.14$, $p = 0.005$ [TGC]; $r = 0.12$, $p = 0.017$ [TRM]). (**B)** The longitudinal analysis using TGC and TRM in Baseline to predict behaviors in Followup2 (PRM percent correct: $r = 0.32$, $p = 0.001$ [TGC]; $r = 0.13$, $p = 0.036$ [TRM]; Attention: $r = 0.13$, $p = 0.012$ [TGC]; $r = 0.19$, $p = 0.006$ [TRM]; AGN negative: $r = 0.19$, $p = 0.001$ [TGC]; $r = 0.15$, $p = 0.001$ [TRM]; AGN positive: $r = 0.19$, $p = 0.001$ [TGC]; $r = 0.12$, $p = 0.003$ [TRM]; SWM between errors: $r = 0.23$, $p = 0.001$ [TGC]; $r = 0.16$, $p = 0.001$ [TRM]; SWM strategy: $r = 0.14$, $p = 0.001$ [TGC]; $r = 0.13$, $p = 0.017$ [TRM]) and in FollowUp3 (TRM: SWM between errors: $r = 0.23$, $p = 0.001$ [TGC]; $r = 0.10$, $p = 0.042$ [TRM], SWM strategy: $r = 0.14$, $p = 0.001$ [TGC]; $r = 0.11$, $p = 0.046$ [TRM]; CGT delay aversion: $r = 0.15$, $p = 0.001$ [TGC]; $r = 0.10$, $p = 0.028$ [TRM]; CGT overall proportion bet: $r = 0.25$, $p = 0.001$ [TGC]; $r = 0.10$, $p = 0.028$ [TRM]; CGT risk adjustment: $r = 0.17$; $p = 0.001$ [TGC]; $r = 0.10$, $p = 0.032$ [TRM]; SDQ: $r = 0.14$, $p = 0.001$ [TGC]; $r = 0.14$, $p = 0.002$ [TRM]).

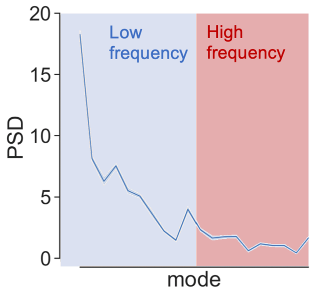


**Figure S10. Energy spectral density of eigenmodes.**

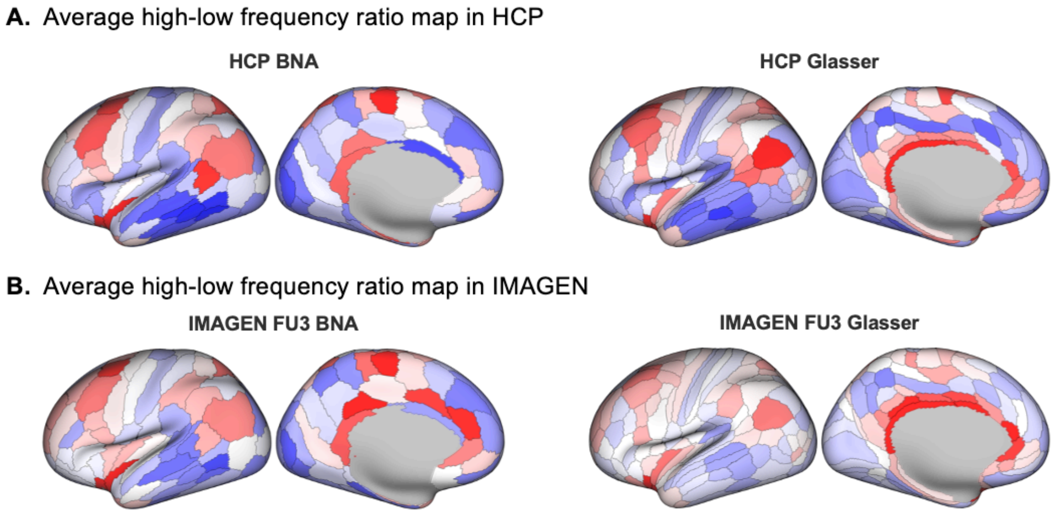


**Figure S11. The replication of average high-low frequency ratio. (A)** The average high-low frequency ratio is stable in different atlases. **(B)** The average high-low frequency ratio map in IMAGEN Followup3 is similar with the ratio map in HCP dataset, both in BNA and Glasser atlas.

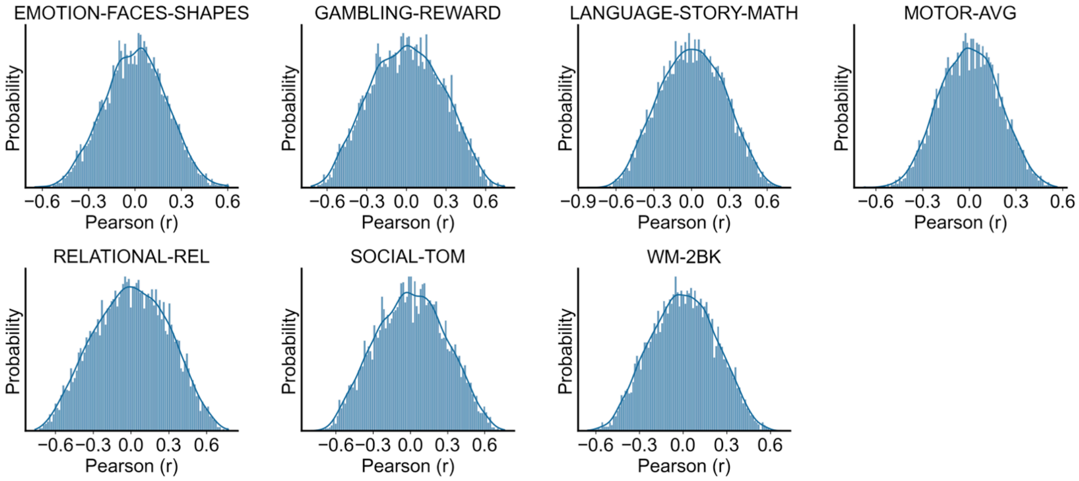


**Figure S12. Correlation between TGC and TGC-AM-correlation.** The distribution of correlation coefficients between TGC and TGC-AM-correlation. Seven contrasts from different behavior domains are shown.

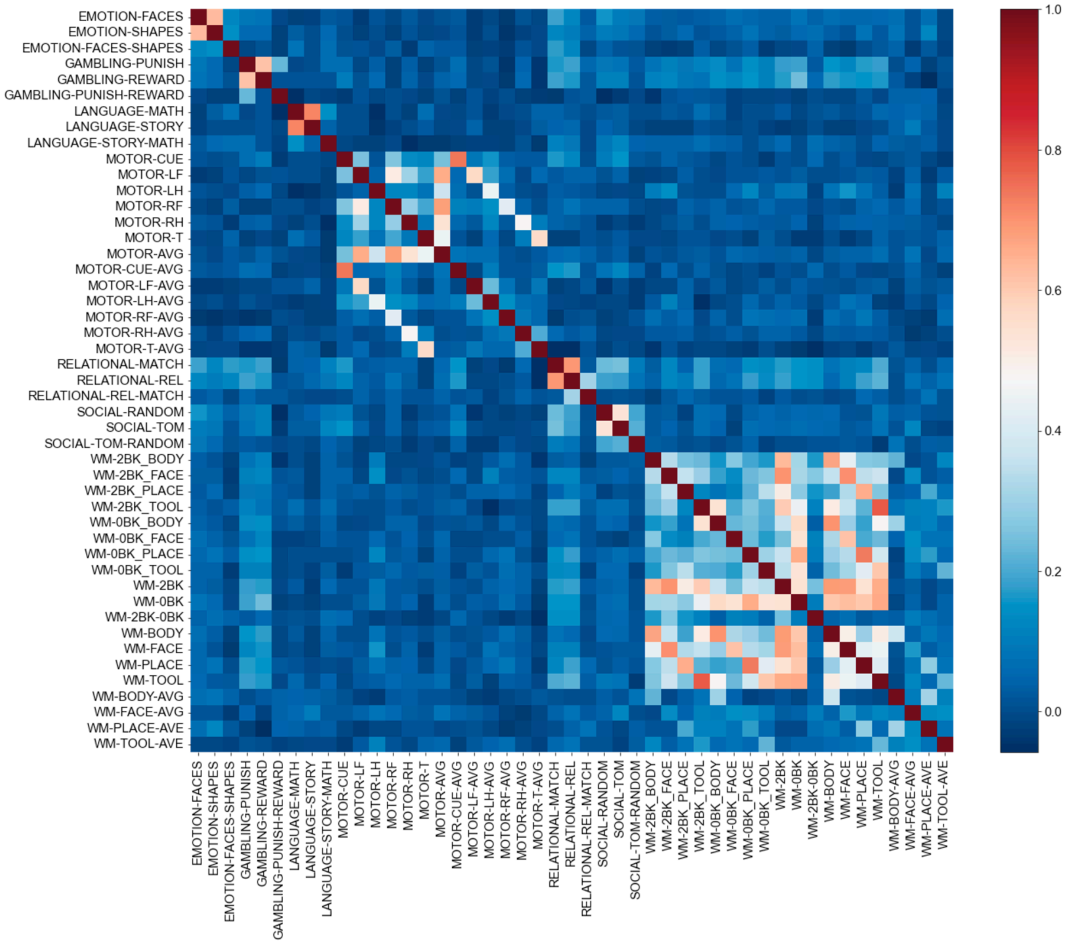


**Figure S13. Correlation between TGC and the number of vertices with significant correlations from 47 contrasts.** For each contrast, we counted the number of vertices showing significant correlations after FDR correction ($p < .05$) for each TGC. We correlated the resultant TGCs between 47 contrasts and found that the similarity between the contrast maps in the same behavioral domain was high.
